# AdaTiSS: A Novel Data-Adaptive Robust Method for Quantifying Tissue Specificity Scores

**DOI:** 10.1101/869404

**Authors:** Meng Wang, Lihua Jiang, Michael P. Snyder

## Abstract

**Motivation:** Accurately detecting tissue specificity (TS) in genes helps researchers understand tissue functions at the molecular level, and further identify disease mechanisms and discover tissue-specific therapeutic targets. The Genotype-Tissue Expression (GTEx) project (Consortium, 2015), and the Human Protein Atlas (HPA) project (Uhlén, et al., 2015) are two publicly available data resources, providing large-scale gene expressions across multiple tissue types. Multiple tissue comparisons, technical background noise and unknown variation factors make it challenging to accurately identify tissue specific gene expressions. Several methods worked on measuring the overall TS in gene expressions and classifying genes into tissue-enrichment categories. There still lacks a robust method to provide quantitative TS scores for each tissue.

**Methods:** We recognized that the key to quantify tissue specific gene expressions is to properly define a concept of expression population. We considered that inside the population, the sample expressions from various tissues are more or less balanced, and the outlier expressions outside the population may indicate tissue specificity. We then formulated the question to robustly estimate the population distribution. In a linear regression problem, we developed a novel data-adaptive robust estimation based on density-power-weight under unknown outlier distribution and non-vanishing outlier proportion (Wang, et al., 2019). In the question of quantifying TS, we focused on the Gaussian-population mixture model. We took into account gene heterogeneities and applied the robust data-adaptive procedure to estimate the population. With the robustly estimated population parameters, we constructed the AdaTiSS algorithm to obtain data-adaptive quantitative TS scores.

**Results:** Our TS scores from the AdaTiSS algorithm achieve the goal that the TS scores are comparable across tissues and also across genes, which standardize gene expressions in terms of TS. Compared to the categorical TS method such as the HPA criterion, our method provides more information on the population fitting, and shows advantages in quantitatively analyzing tissue specific functions, making the biology functional analysis more precise. We also discuss some limitations and possible future work.

## 1 Introduction

In the analysis of gene expressions, one important task is to detect tissue specificity (TS) of genes, i.e., whether the genes are housekeeping, or are significantly differentially expressed in one or multiple of tissues. Accurately detecting TS in genes helps researchers understand tissue functions at the molecular level. Understanding TS at the molecular level helps us further identify disease mechanisms and discover tissue-specific therapeutic targets (Greller and Tobin, 1999; Kim, et al., 2017).

The development of high-throughput technologies greatly improves the study of tissue specificity in genes. The Genotype-Tissue Expression (GTEx) project (Consortium, 2015) is an ongoing effort to build a comprehensive public resource to study tissue-specific gene expression and regulation. Based on the RNA-sequencing (RNA-seq) technology, up to version 7, the GTEx project has generated RNA expressions for 18,777 human protein-coding genes from 11,688 samples and 53 tissues. Other gene expression databases include the Tissue-specific Gene Expression and Regulation (TiGER) (Liu, et al., 2008), the Human Protein Atlas portal (HPA) (Uhlén, et al., 2015), and the Tissue-specific Gene DataBase in cancer (TissGDB) (Kim, et al., 2017).

Large-scale databases provide valuable resources but also make the task to quantify TS more challenging. If there were only two tissues, we could apply the *t*-test testing for differential expressions. But now we have more than 50 tissue types in the GTEx project. Pairwise or triplet comparison is computationally inefficient and unfeasible, and creates a burden in multiple hypotheses testing (Cavalli, et al., 2011). Moreover, sample variation and background noise make it harder to identify truly high-expressed samples. We expect that a good TS measurement should have the properties not only identifying overall tissue specificity to tell house-keeping genes and tissue-specific genes, but also sensitively and robustly measuring expression specificity for each tissue.

The previous work (Kryuchkova-Mostacci and Robinson-Rechavi, 2017) reviewed several metrics measuring TS but mainly focused on the methods of measuring overall specificity, not identifying the specific tissues. Categorizing a gene to be tissue-specific or ubiquitous only based on the overall TS score may mask potential subcategories such as genes specific in multiple tissues. There are a few methods for measuring TS in the tissue level including z-score (Cheadle, et al., 2003), the preferential expression measure (PEM) (Huminiecki, et al., 2003) the expression enrichment (EE) (Yu, et al., 2006), and the specificity measurement (SPM) (Xiao, et al., 2010). The last three measurements for TS rely on the constraint of total sum (or *l*_2_ norm) of tissue expressions. With or without an extremely high expression, the TS measurements can be dramatically changed.

The z-score relaxes such constraint and is easy to interpret, usually applied in analyzing only a few tissues. In the large-scale data analysis, when we measure many tissues, some tissues having related functions could have similar high expressions. These outliers can be in unexpectedly large proportion. If we ignore the effect from outliers and simply estimate the mean and variance from all the samples, the TS based on the traditional z-score can be too conservative.

(Kadota, et al., 2006) realized the influence from the high or low expressed outliers and considered all the combinations of inlier and outlier tissues in the method ROKU. The drawback of their method is that it cannot detect the case where there is no outlier tissue. (Zhang, et al., 2017) took more care on the outlier expression in the method of the specifically expressed gene detection tool (SEGtool). They first defined the outlier expression region and applied computational clustering tools to categorize each tissue as low, median or high expression. Since those expression categories depend on the initially defined outlier region, robustness of the outlier region affects the stability of their categories. (Uhlén, et al., 2015) took a direct way to cluster inlier and outlier tissues based on the fold change in the HPA project. Their HPA criterion differentiated the highly expressed tissues (outliers, with at least 5 fold-change) from the rest of the tissues (inliers), in the comparison of tissue expressions in terms of the transcripts per kilobase million (TPM) or the reads per kilobase million (RPKM) from RNA-seq. Based on the inlier and outlier configuration, they classified genes into six categories: tissue enriched (one outlier); group enriched (multiple outliers); tissue enhanced; expressed in all; mixed; and not detected. Although their HPA criterion provides such fine categories, it still cannot quantify the TS of the enriched tissue (highly expressed tissue), nor the TS profiling of all the tissues. For a particular gene, the highest tissue might be expressed as a 10 fold-change or even a 20 fold-change compared to the rest of the tissues, but it is only characterized as “enriched” without a quantified score. More importantly, the specificity based on the fold-change is not comparable across genes. For one gene, an enriched tissue having a 10-fold change compared to the rest does not mean that tissue is more specific in another gene having a 5 fold-change compared to the rest. These categorizing methods motivate us to develop a robust method to quantify gene expressions in terms of tissue specificity scores. With the well-estimated robust TS scores, we can quantify specificity for each tissue in each gene and compare TS across genes, which builds standardized scores for future comparison across -omics. Such standardized scores also provide quantitative TS comparison in GO terms and pathways, refining the biology functional analysis to a precise level.

The outline of the rest of this paper is as follows. In **Section 2**, we introduce the key concept of population in the problem of defining TS, then propose a data-adaptive and robust estimation method to get TS scores (AdaTiSS) under Gaussian population, and we provide the algorithm to implement it. We put simulation studies with method comparisons and the development of data-adaptive estimation procedure under *t*-population in the **Supplementary Materials**. In **Section 3**, we study the performance of AdaTiSS and the HPA criterion in the real data from the GTEx project, comparing the performance in detecting housekeeping gene candidates and tissue-specific genes. Finally, in **Section 4**, we conclude the paper, and discuss some limitations and future work.

## 2 Methods

### 2.1 Population

From previous works, we can see the first task to quantify TS is to distinguish inlier and outlier tissues, where the outliers can be the tissue-specific expressions. The outlier expressions vary from gene to gene. Some genes are highly expressed (enriched) in only one tissue, while some are highly expressed in multiple tissues. Among enriched tissues, some expression is moderately high and some is extremely high. Due to the complexity of the tissue-specific outliers, our contribution is a focus on the inliers, defining the concept of expression *population*. Inside the population, some tissue is highly expressed while some is not, but there is still a balance. Meanwhile, the tissue-specific expressions that go outside such balance become outliers. In this work, we focus mainly on the outlier in high expression but not in low expression.

After introducing the concept of population, the next question is what variations are inside the population. Most of the previous works summarized the tissue information in medians, then defined TS by comparing tissue medians, which only considered the variation between tissues. We consider our population containing not only between-tissue variation but also within-tissue variation. We allow biological replicates in each tissue to incorporate within-tissue variation. When comparing sample expressions from multiple tissues, we consider the main effect in the population to be the tissue effect, which is confirmed in several studies from the analysis of tissue sample clustering (Jiang, et al., 2019; Melé, et al., 2015). In our own work of (Jiang, et al., 2019), we provided a website resource TSomics (http://snyderome.stanford.edu/TSomics.html), presenting RNA and protein expressions for all the quantifiied human protein-coding genes. From there, for the majority of genes, the abundance distribution across tissues appears as one big density bump for the main population with some specific tissues outside the population. In biology, it is conventional to take the logarithm transformation assuming Gaussian noise in the analysis of expression data (Hill, et al., 2008; Melé, et al., 2015). Based on our experience and other previous works, here we consider the population distribution as Gaussian but we do not an assume outlier distribution.

Once we can identify such Gaussian population, it is natural to quantify TS in terms of the robust z-score:

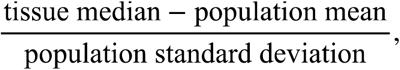

measuring the distance from the tissue median to the population mean, standardized by the population standard deviation for each tissue. In contrast to the traditional z-score, the mean and variance in our robust z-score are from the population and thus the score is more sensitive to the TS outliers. Importantly, when such tissue balances are similar in all the genes, then the robust z-scores are comparable across genes.

The remaining question is how to estimate the population given the presence of various outlier expressions. In statistics, we formulate this question as robust estimation for the population in a mixture model. The literature of robust estimation is very rich, such as median and the median absolute deviation (MAD), Efron’s method of empirical null distribution (Efron, 2005), and Tukey’s biweight fitting, etc. In the cases where all the samples can be modeled as a mixture model of multiple Gaussian components, one can use the Bayesian method from the EM algorithm (Cavalli, et al., 2011). They built a pipeline in the method of SpeCond to determine which Gaussian component belongs to the population, and which to the outliers.

If the outliers are in a large proportion, and if their distribution is complicated and unknown, the problem becomes challenging. We still lack a robust, automatically data-adaptive method to estimate the population information. Recently in a general linear regression problem, we developed a novel data-adaptive robust estimation based on density-power-weight under unknown outlier distribution and non-vanishing outlier proportion (Wang, et al., 2019). In the question of quantifying TS, we restrict the multivariable model analyzed in (Wang, et al., 2019) to a univariate model in the Gaussian population, and robustly estimate population information to get our data-adaptive quantitative TS in the form of robust z-score (AdaTiSS). Our AdaTiSS takes into account gene heterogeneities under various outlier proportions and magnitudes. It achieves robustness and data-adaptiveness by selecting a turning parameter based on the data to optimize the population estimation. We summarize the procedure and the algorithm in the following subsections, and more statistical analysis can be found in (Wang, et al., 2019).

### 2.2 Data-adaptive robust TS scores

Consider a gene expression matrix *X* for *n* genes from *m* samples. Each row contains the expressions from *T* tissues for one gene and each tissue can have several biological replicates. Column expressions are from one sample across all the genes. When the samples are collected from different tissues, we consider the tissue effect is the major factor. Under the log scale, we model the majority of the samples coming from Gaussian population distribution to capture the tissue main effect. The outlier samples may reflect tissue specificity. Each expression in one gene has some probability from the population distribution or from the outlier distribution. To fix the idea, we formulate the distribution of the gene expressions for each gene as a Gaussian-null mixture model,

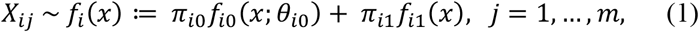

where *f*_*i*_ is the mixture density for gene *i*, *f*_*i*0_ is the Gaussian population (the null) density parameterized by 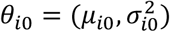 with mean *μ*_*i*0_ and variance 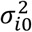, *f*_*i*1_ is the unknown outlier density, π_*i*0_ ∈ (0.5, 1] is the population proportion, and π_*i*1_ = 1 − π_*i*0_ is the outlier proportion. Our model allows heterogeneous genes having population distributions with different (π_*i*0_, *θ*_*i*0_)’s. In contrast to a traditional method assuming the full model on all the samples, our method relaxes the assumption on the outlier distribution. Here we do not assume that π_*i*1_ vanishes to zero as increasing the sample size. In our task to quantify TS, the outlier proportion may not be small, especially for a gene highly expressed in multiple tissues. Under the unknown outlier distribution and non-vanishing outlier proportion, we would like to estimate the null parameters (π_*i*0_, *θ*_*i*0_) for each gene.

Applying the method in (Wang, et al., 2019) to estimate the null parameters, we take the technique of weighting the mixture density by the density power 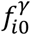,where *γ* ≥ 0, to down weight the outliers and therefore purify the reweighted mixture model. The rational is that under a proper *γ*, the outliers go to the tails of the reweighted null density and thus the outliers do not contribute much to the population estimation. The technique of density power weight played an important role in robust estimation (Basu, et al., 1998; Fujisawa, 2013; Fujisawa and Eguchi, 2008; Windham, 1995). In our setting, we construct a robust criterion from weighted loglikelihood function under the system parameter *γ* ≥ 0,

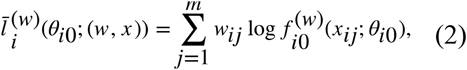

where

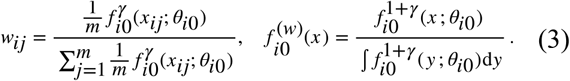

Taking *γ* = 0, our weighted log-likelihood criterion is the average of ordinary log-likelihood functions. Our robust estimate is

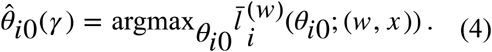

For the Gaussian-null mixture model, the estimates for the null parameters are summarized in the **Proposition 1**. The estimate for *θ*_*i*0_ is the same as based on *γ*-cross entropy in (Fujisawa and Eguchi, 2008) and the estimating equation in (Windham, 1995). Our estimator for π_*i*0_ agrees with the result from minimizing the density power score in (Kanamori and Fujisawa, 2015). The estimation can be done in an iterative fashion. We illustrate the iterations of the population fitting in **Figure 1**. We can see, along the iterations, the estimated population densities are approaching to the underlying population density.

**Figure 1:**
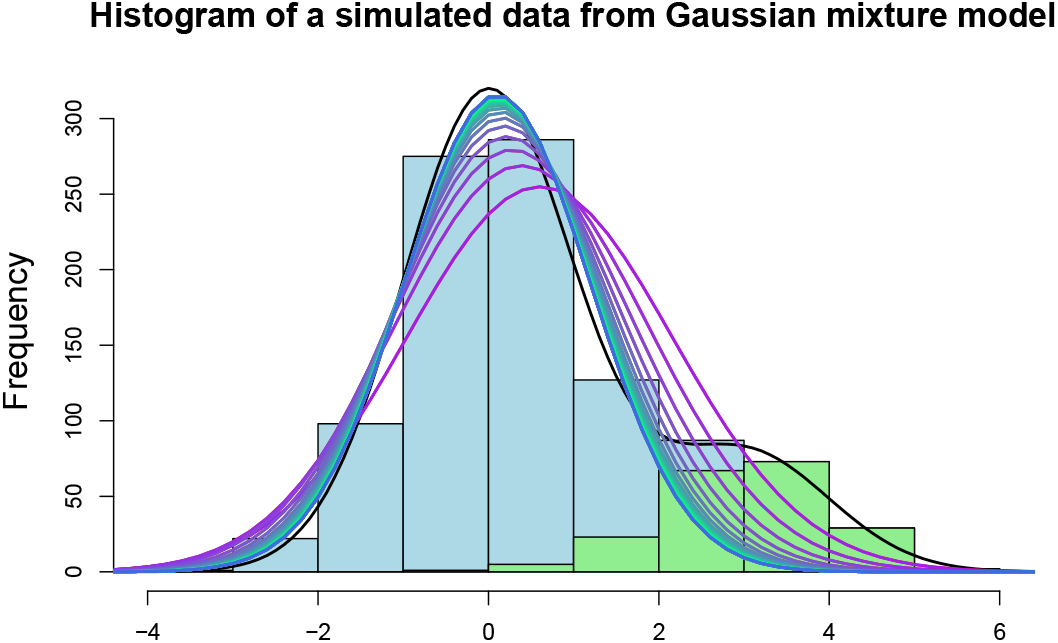
Histogram of a simulated data from Gaussian mixture model, i.e., *X*_*i*_ ~*iid* 0.8*N*(0,1) + 0.1*N*(3,1), *i* = 1,…, 1000. The blue bars indicate the samples from the population distribution and the green ones for the outliers. The black curve is the underlying mixture density. The curves in gradient colors from purple to blue are the fitted population densities along iterations and the blue curve is the final fitted density.

#### Proposition 1.

*For gene i, consider the expressions* 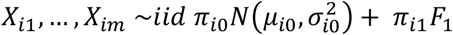, *where F*_*1*_ *is the unknown outlier distribution. Our robust weighted log-likelihood function is*

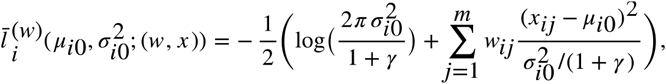

*The robust estimates for* 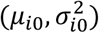 *and the estimate for π*_*i*0_ *satisfy*

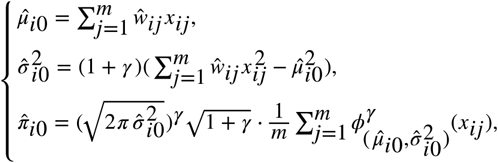

*where* 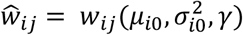 *is defined in (3)*.

So far, the robust estimation is implemented under a predefined system parameter *γ*. As pointed in many previous papers (Windham, 1995, Fujisawa and Eguchi, 2008), the *γ* balances robustness and efficiency of the estimation. **Figure 2** shows a simulation result on fitting the population density under various *γ*’s. When *γ* = 0, 0.5, the fitting is affected by the highly expressed outliers. When *γ* = 3, the fitting becomes locally trapped such that it only locally fits well on a small set of null points. From the density fittings shown in **Figure 2**, what really matters is the goodness-of-fit (GOF) of the fitted population density to the mixture density on the population points. Hence, we evaluate the GOF in the measure of population distribution by

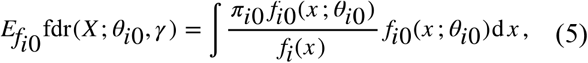

where *fdr*(*x*) ≔ *P*(*x is from the population |x*) = π_0_*f*_0_(*x*)/*f*(*x*) is the local false discovery rate introduced in (Efron, 2005). Comparing the estimated (5) to an approximate value one gives our selection criterion for *γ*,

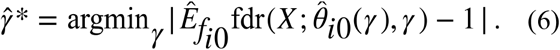

**Figure 2:**
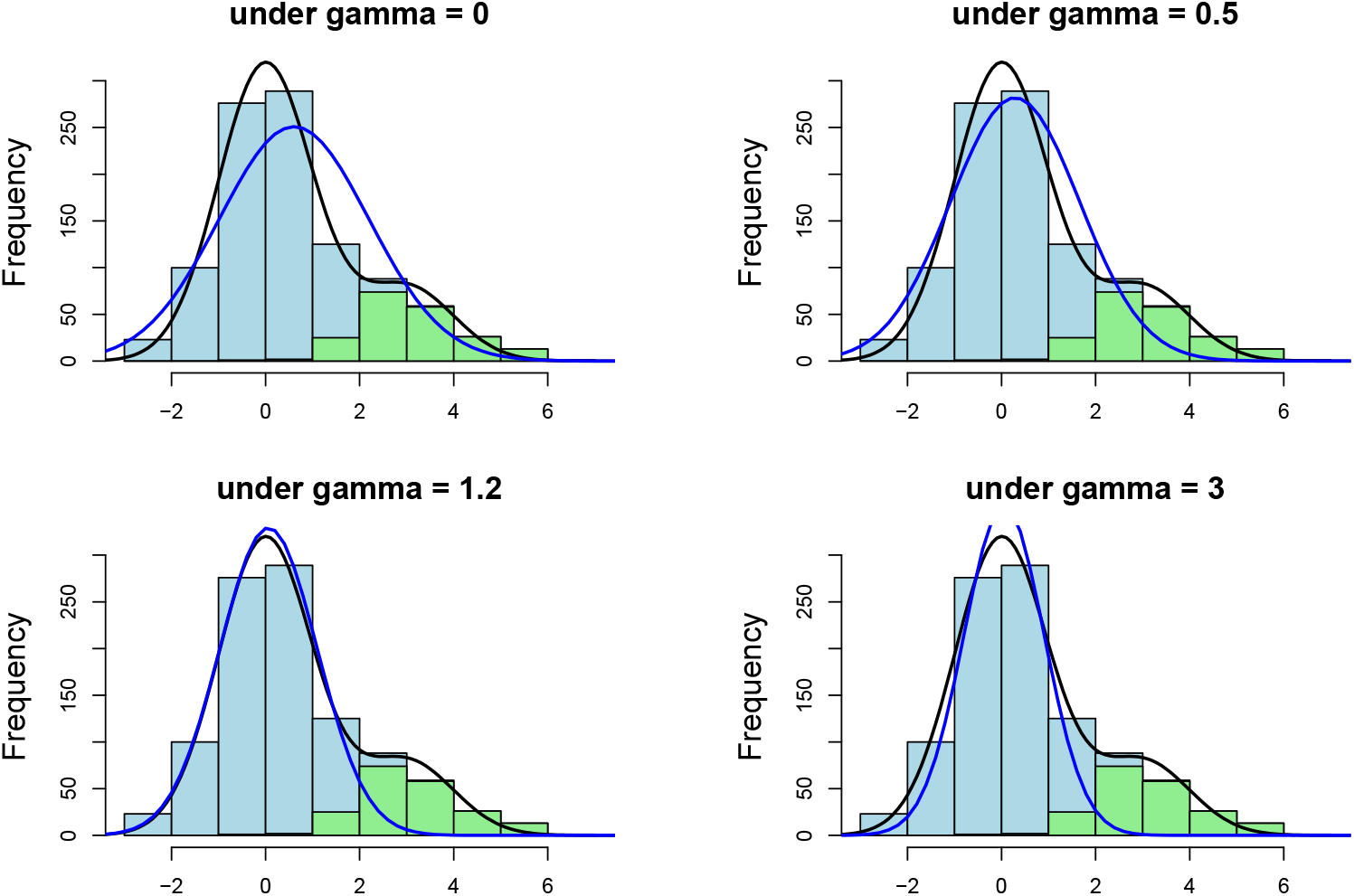
Histogram of 1000 X’s from 0.8 *N*(0,1) + 0.2*N*(3,1) in the simulations where the points from N(0,1) are colored in blue and the ones from N(3,1) in green. The the black curve is the underlying density for the mixture model. The blue curve is the fitted partial density of 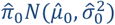 under *γ* = 0, 0.5, 1.2, 3, where *γ* = 1.2 is selected from our data adaptive selection procedure.

Our data-adaptive robust estimation is the 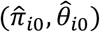 under the selected 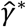 from (6). The selected 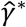 varies from gene to gene, trying to optimize the estimation for each gene. (Wang, et al., 2019) provided more statistical analysis on the data-adaptive procedure. Here, we apply this new procedure to estimate the population distribution under the Gaussian-null mixture model formulated in (1) in the problem of quantifying TS.

We compare our data-adaptive robust population estimation to several methods in simulations, and evaluate their performances on estimating the population information in terms of mean squared error (MSE). The methods under comparison include the fixed *γ* robust estimation, Tukey’s biweight estimation, the EM algorithm under two Gaussian mixtures, and Efron’s local fdr estimation. In the simulation studies, we consider the outlier distribution to be Gaussian or *t*-distribution under various model parameters. The comparison results are summarized in the **Supplemental Materials**. We conclude that the *γ*-robust estimation methods perform better than the other methods in various situations. The EM algorithm only works well when the mixtures are well separated. The local fdr method is mainly designed under light outliers. Our data-adaptive estimation performs better overall than the fixed *γ* estimation, especially under the outliers are in a large proportion. It is more adaptive to various outlier distributions.

In the **Supplementary materials**, following the similar analysis in Gaussian-population mixture model, we develop a data-adaptive procedure for the *t*-population mixture model. We find our data-adaptive procedure prefers light tailed population density. In the real practice when applying our data-adaptive procedure, we suggest taking log transformation or other transformation to make the data more Gaussian distributed in order to maintain the detection power of our procedure.

Based on our data-adaptive robust estimation for the population distribution, we establish sample outlying *z*-score under the selected 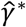 for gene *i* in sample *j* as

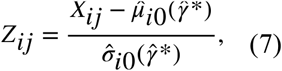

and tissue level specificity z-score for gene *i* in tissue *t* as

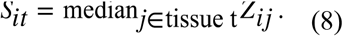

The TS score *S* gives a quantitative measurement on the specificity of a tissue expression relative to the sample population for each gene. This makes it possible to compare tissue specificities across tissues and also across genes.

From our quantitative TS scores, we can also define tissue enrichment categories. Define a gene to be enriched if it includes at least one highly expressed tissue and the highly expressed tissues are in the top *t*\* highest scores, where *t*\* = *argmin*_*t*_ {*S*_*i*(*t*)_ − *S*_*i*(*t*+1)_ ≥ 1.5, *S*_*i*(*t*)_ ≥ 3} and *S*_*i*(*t*)_′*s* are the ordered tissue scores sorted in decreasing order in one gene. The 1.5 z-score gap is analogous to the concept of a 5 fold-change gap between the enriched tissues and the rests in the definition of the HPA categorizing method.

### 2.3 Algorithm of AdaTiSS

In this section, we develop the algorithm to get our data-adaptive TS scores (AdaTiSS). For the intensity data from the microarray or the mass spectrometry platform, the Gaussian distribution is usually used as a convention after taking the log transformation. One can directly apply our robust fitting method there. For the data from RNA-seq, one can work on the log transformed standardized TPM or RKPM. When comparing multiple tissues, it could happen that in some low expressed or unexpressed tissues, their TPM (or the RKPM) are zeroes. To deal with zeroes when taking the log transformation, there are several ways. One way is to add a small amount of positive perturbation on the TPM near zero. Another way is to ignore the expression differences when the TPM is less than one. Here, we take the approach of taking TPM less than one to be one. Since there could be a density peak at log(1), we make adjustments on defining the population and provide criteria to determine whether such zero peak contributes to the population. Another possible approach dealing with the zero expressions is to take square root or cubic root, then the variation at zero is under controlled. Since here we more focus on the high expressions, we prefer the log transformation to more control the variation in high expressions. We detail the procedures to implement AdaTiSS below.

**Step 1: Preprocessing.** First, to do normalization or standardization and to remove batch effects, as conventions. Since this paper does not extend the discussion on the normalization, we assume the data is well normalized or standardized before applying AdaTiSS. We allow biological replicates in the data. For the data having technical replicates, we take the average expressions within the tissue from the same subject. We filter the genes with low expressions in all the tissues.

**Step 2: Fitting the population distribution.** Following the same notations in the previous subsection, we estimate population mean, standard deviation and proportion from **Algorithm 1**. For the TPM or RPKM data, we take all the negative expressions in the log scale concentrated at zero, then fit the Gaussian population on all the positive expressions. We then adjust the overall population in the next step.

**Step 3: Calculating TS scores**. For the log transformed intensity expressions, we obtain their TS scores based on the equations (7)-(8) with robustly fitted 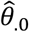 from Step 2. For the log transformed TPM or RPKM expressions, we make adjustments in two complementary cases to determine the overall population. Genes in Case (i): their fitted 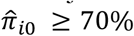 from Step 2 and the zero peak lies outside three standard deviations (SD) away from the fitted mean, as an example shown in the top row in **Figure 3**. For Case (i), we take the overall population as the fitted Gaussian bump from the positive expressions. Genes in Case (ii): their fitted 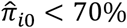 or the zero peak lies inside three SDs away from the fitted mean, which is the complement to Case (i), as an example shown in the bottom row in **Figure 3**. In Case (ii), we take the samples inside the ± 3SDs away from the fitted mean and also include the zeroes as the overall population. Then the population information is obtained by taking the mean, standard deviation, and the proportion of these inliers.

**Figure 3:**
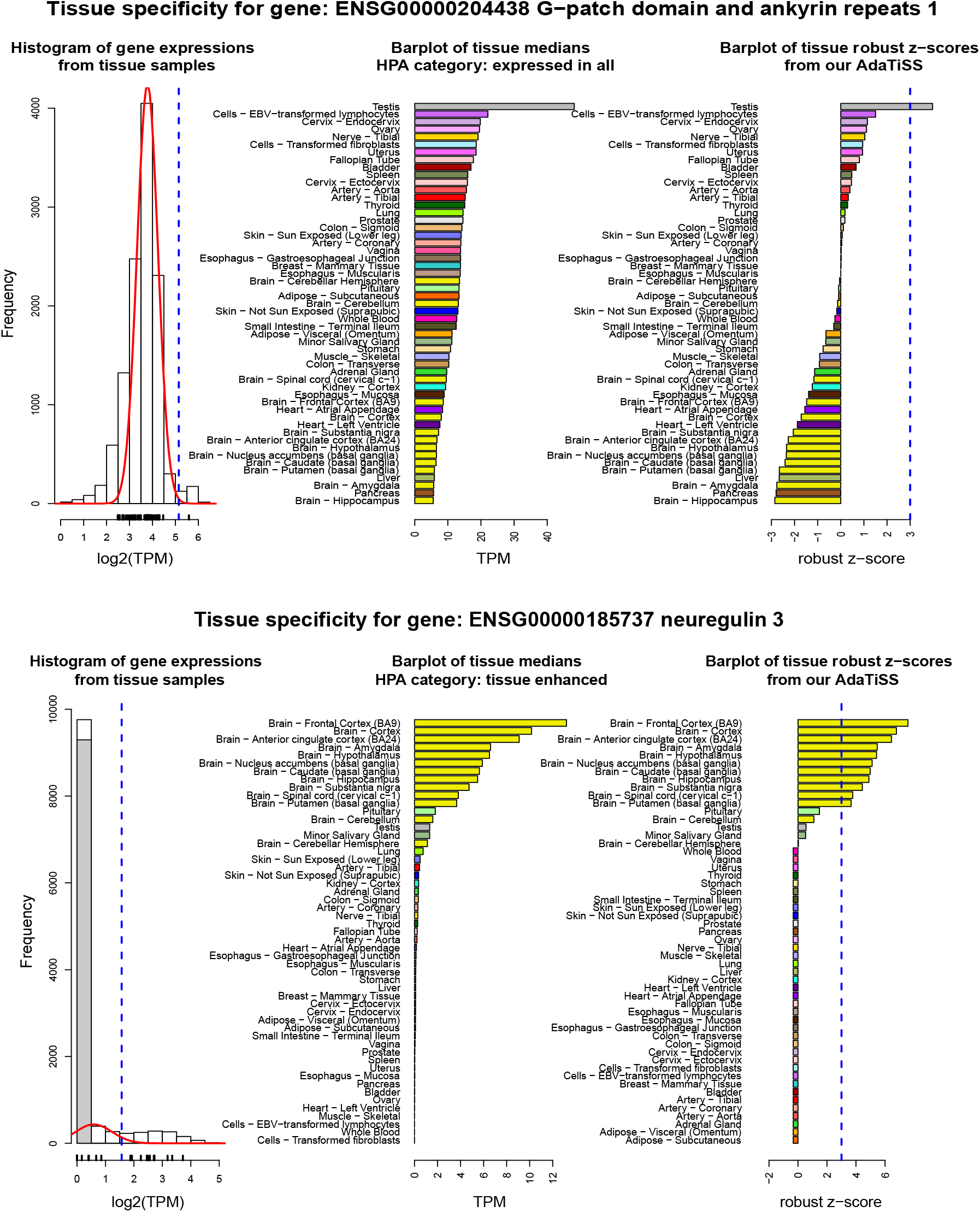
Two example of tissue specific genes. The plots in the top row are for a testis-specific gene (GPANK1) in Case (i) defined in Step 3 and the one in the bottom row are for a brain-group-specific gene (NRG3) in Case (ii). The histograms in the first column are from gene sample expressions in log(TPM) from GTEx dataset in version 7. In the histograms, the curves are proportional to the robustly fitted density of 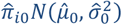; the vertical dashed line is the threshold as three standard deviations from the fitted mean in the right tail; the small bars at the bottom indicate tissue medians; the grey partial colored bar in the left tail are for the zero log(TPM). The plots in the middle column are the bar plots of tissue median log(TPM) and indicated by the HPA category. The plots in the right column are the bar plots of our TS scores. The dashed lines are at 3 standard deviation from the fitted mean.

**Step 4: Diagnosis.** Since the real data may be more complicated, we add one diagnosis to check whether the fitting population proportion is small, trying to indicate the case of the fitting being locally trapped. If the fitted population proportion is less than 70%, we mark that gene and further check its fitting plot.

#### Algorithm 1: Data-adaptive robust estimation for the population parameters

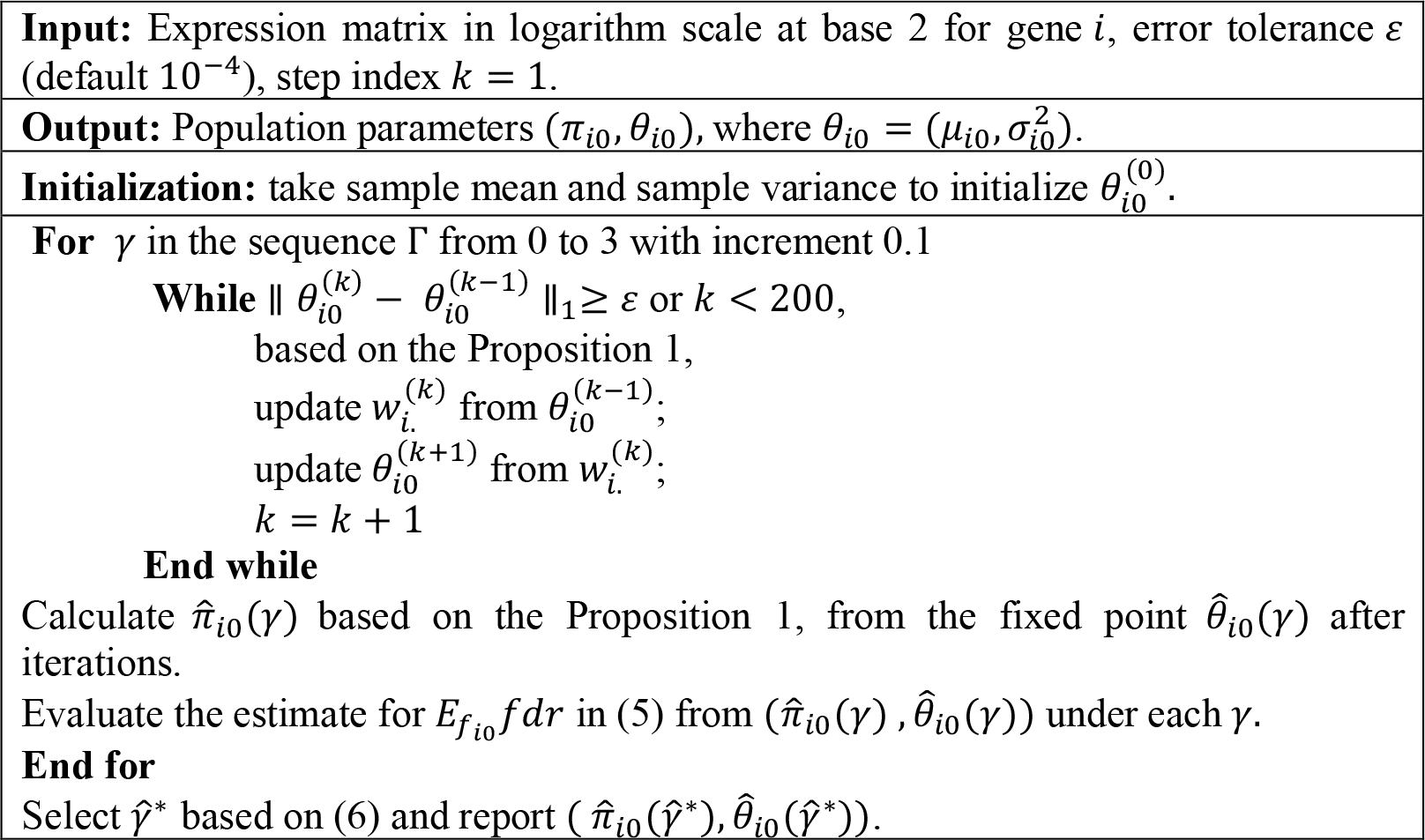

## 3 Results

From the simulation studies our data-adaptive quantitative TS method (AdaTiSS) is more adaptive to heterogenous cases than other methods in the comparison. The HPA categorizing method has been applicated to discovery several biological findings (Uhlén, et al., 2015). In this subsection, we investigate the comparability of our TS scores to the HPA categories on the gene expressions from RNA-seq in the GTEx project (the dataset is from the release in version 7, downloaded from http://www.gtexportal.org). The HPA criterion defines genes into six categories: tissue enriched (one tissue is at least five fold-change higher than the rest of the tissues), group tissues (multiple tissues are all at least five fold-change higher than the rest of the tissues), tissue enhanced (at least one tissue is at least five fold-change higher than the average expression of all the tissues), expressed in all (the gene is detected in all tissues), mixed, and not detected. The GTEx project has quantified RNA expressions in 18,777 protein-coding genes from 11,688 samples and 53 tissues, up to the version 7. The tissue level information is summarized as median TPM of tissue samples. After removing the genes in which all the tissue medians less than one, we take the rest 17,719 genes for our comparison analysis.

### 3.1 Housekeeping genes

The housekeeping genes are supposed to be stable, uniformly expressed in all the tissues, and maintain the basal cellular functions (Eisenberg and Levanon, 2013). From RNA expressions, we compare the lists of the housekeeping genes based on our criterion from the results of AdaTiSS and the HPA criterion. Since the housekeeping genes should be expressed in all the tissues, we filter out the low expressed genes and end up 7,938 genes in which all tissue medians are greater than or equal to 1 TPM. Among these genes, the HPA criterion categorizes 7,054 genes to be “expressed in all" as the housekeeping gene candidates. In contrast to the HPA criterion which only gives a category, we further give the population information for the candidates based on our robust fitting. **Figure 4** shows that some candidate genes from the HPA criterion still have large variations and relatively low population proportions. Adding the population information, we get our criterion for the housekeeping genes requiring (1) the population proportion ≥ 80%, (2) the population fitted robust standard deviation ≤ 1, and (3) *max*_*t*_|*S*_.*t*_| ≤ 3, where *S* is defined in (8) in **Section 2**. We finally get 2,056 housekeeping gene candidates, where 2,040 genes are in the “expressed in all” category and 16 genes in “tissue enhanced” from the HPA criterion. We perform the GO terms enrichment analysis in the biological process on the two housekeeping gene lists from the tool of “GOrilla” (Eden, et al., 2009). We then reduce the redundancy among the significant GO terms by the method of “REVIGO” (Supek, et al., 2011). For the reduced-redundancy set of GO terms with the frequency ≥ 40% (the frequency of the GO term in the GOA database), the two lists have 70% agreement in 7 terms including: metabolic process, cellular metabolic process, nitrogen compound metabolic process, cellular macromolecule metabolic process, macromolecule metabolic process, organic substance metabolic process, and primary metabolic process. The list from the HPA criterion has three more terms: biological process, cellular process, and cellular response to stimulus. For the redundancy-reduced set of GO terms with the frequency < 40%, two lists have 16% agreement in 54 terms and the list from the HPA criterion has 286 more. We can see that the list from the HPA criterion tends to capture the detailed specific functions, while added the population and the TS scores constraints, the list from our criterion mainly focuses on the general functions, which more reflects the property of the housekeeping genes maintaining basic cellular functions. The list of the redundancy-reduced significant GO terms in biological process from our criterion is shown in **Figure 5.**

**Figure 4:**
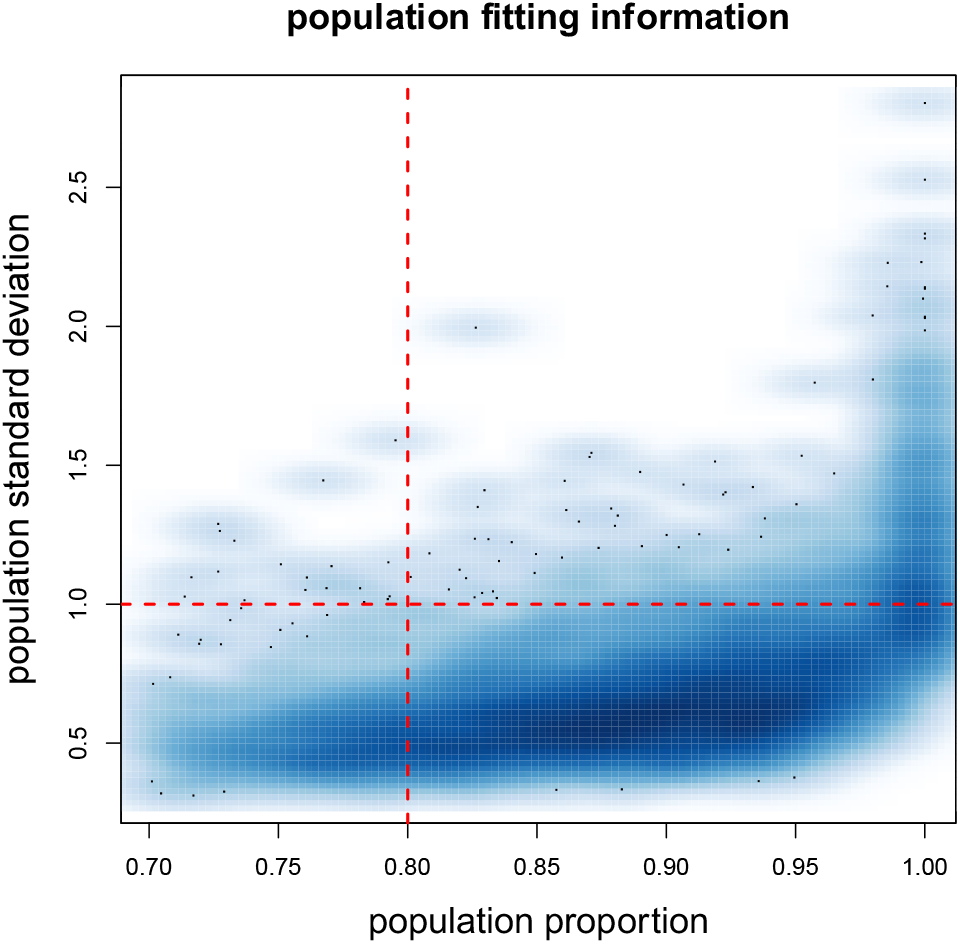
Scatter plot of the population proportion versus population standard deviation (SD) from our robust fitting for the genes in “expressed in all” category from the HPA criterion. The horizontal line is at SD =1 and the vertical line is at proportion =0.8.

**Figure 5:**
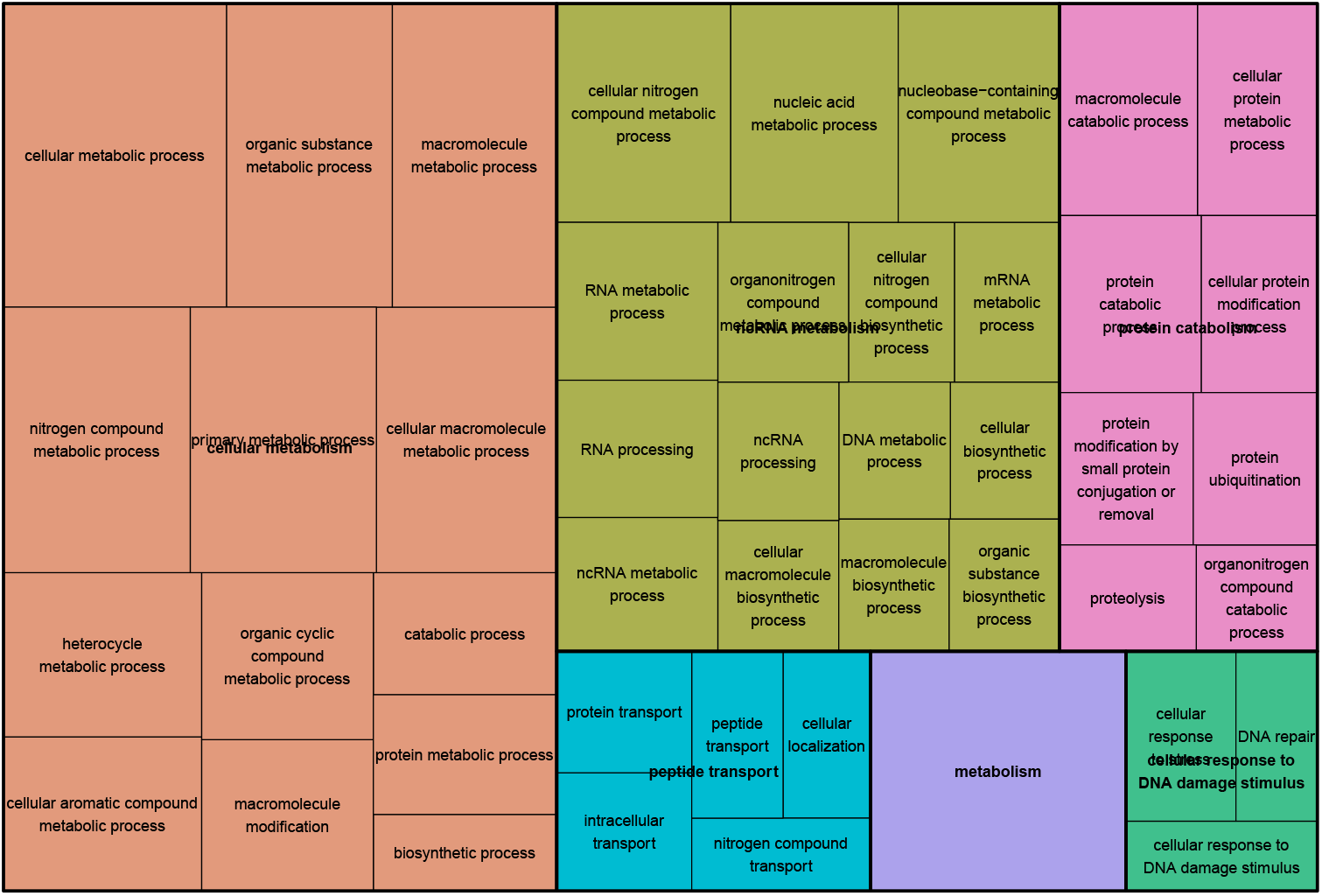
REVIGO Gene Ontology treemap (Supek, et al., 2011) of housekeeping gene candidates from our criterion added constraints on the population information and the TS scores.

### 3.2 Tissue specific genes

In the example of gene GPANK1 shown in the top row of **Figure 3**, the HPA criterion categorizes this gene to be “expression in all”, while the score of testis from our AdaTiSS is above 3 indicating the gene enrichment in testis. Another example is from gene NRG3 shown in the bottom row of **Figure 3**. Its HPA category is “tissue enhanced”, while our TS scores show that this gene is highly expressed in a group of brain regions. These examples show that the HPA categorizing method may miss sub-category for the tissue-specific gene expressions. This is where our quantitative scores come in.

To categorize the genes with highly expressed tissues, the HPA criterion reports 1,879 tissue enriched genes, 2,078 group enriched genes, and 4,350 tissue enhanced genes. By the definition of a tissue-enriched gene in **Section 2**, we identify 4,356 tissue-enriched genes, where 1,821 genes are in the HPA category as “tissue enriched”, covering 97% of the total tissue-enriched genes from the HPA criterion. Among our tissue-enriched genes, there are 1,221 genes in the HPA category as “group enriched”, 1,011 genes as “tissue enhanced”, 253 genes as “expressed in all”, and 50 genes as “mixed”. The two criteria have similar ability to identify single-tissue-enriched genes. To identify multiple-tissue-enriched genes, due to different thresholds, there could be different in categories. Only assigning a category may not be enough to characterize a more complex TS profiling. Our AdaTiSS feeds the need of profiling quantitative tissue specificity scores for all the tissues.

Taking testis-enriched genes as an example, we compare the gene categories from our criterion to the HPA criterion, then investigate their biological functions. From our TS scores, we identify in total 1,866 testis-enriched genes. The histogram of the TS scores in testis-enriched genes is shown in **Figure 6**, colored by the HPA categories. We can see that the genes with high testis scores are also in the “tissue enriched” category from the HPA criterion. Based on the quantitative scores, we can further quantify TS scores in the level of gene sets such as the GO terms, by taking the averaging of our TS scores for each tissue across the genes belonging to the same GO term. **Figure 7** shows the scatter plot of the TS scores in the GO term level versus the minus of the log of adjusted *p*-values based on the BH procedure (Benjamini and Hochberg, 1995). The plot is referred to the “GOplot” package (Walter, et al., 2015). The significant GO terms also have high TS scores. The right panel in **Figure 7** lists the topmost significant GO terms including: meiosis, gamete generation, spermatogenesis et. al, which represent testis specific functions.

**Figure 6:**
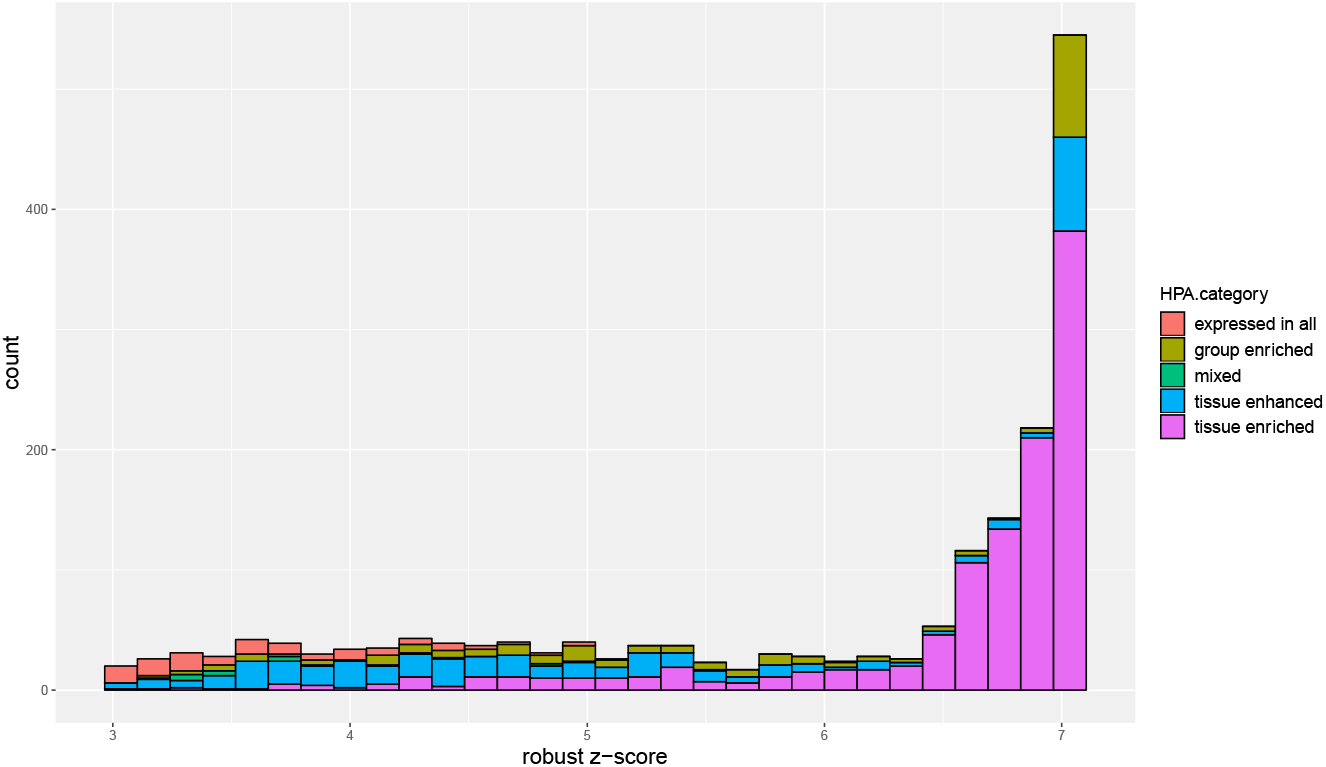
Histogram of testis our TS robust scores from the testis-enriched genes, colored by the HPA categories.

**Figure 7:**
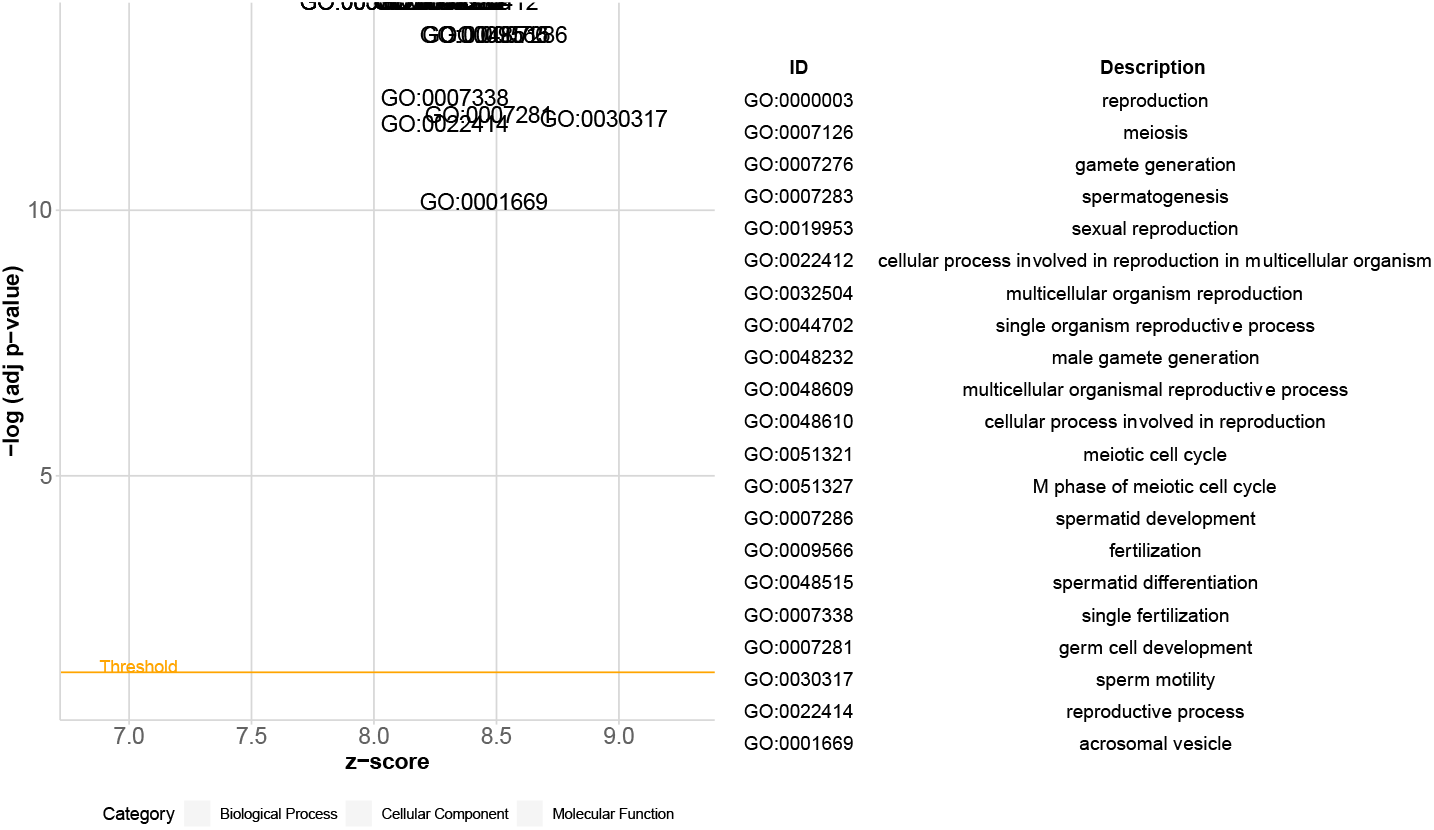
Bubble plot of the average of our TS robust z-scores in testis versus the minus of log_10(p-value) in GOterms from the testis-enriched genes. The table in the right panel shows the topmost significant GO terms. The bubble plot is referred to the package GOplot (Walter, et al., 2015).

Moreover, our robust quantitative TS scores make it possible to compare biological function more precisely. For example, consider the TS scores across 53 tissues for the genes belonging to the GO term: neurological system process, shown in **Figure 8**. We can see that there are a bunch of genes highly expressed in the brain regions, as expected. We further consider the children GO terms of the neurological system process term based on the hierarchical structure of the GO terms. **Figure 9** shows the TS scores across different brain regions for those children GO terms. The results confirm our basic understanding such as the neuronal action potential propagation gets more involved with brain cerebellum and cortex, and the detection of mechanical stimulus involved in sensory perception of sound more involved with cerebellum.

**Figure 8:**
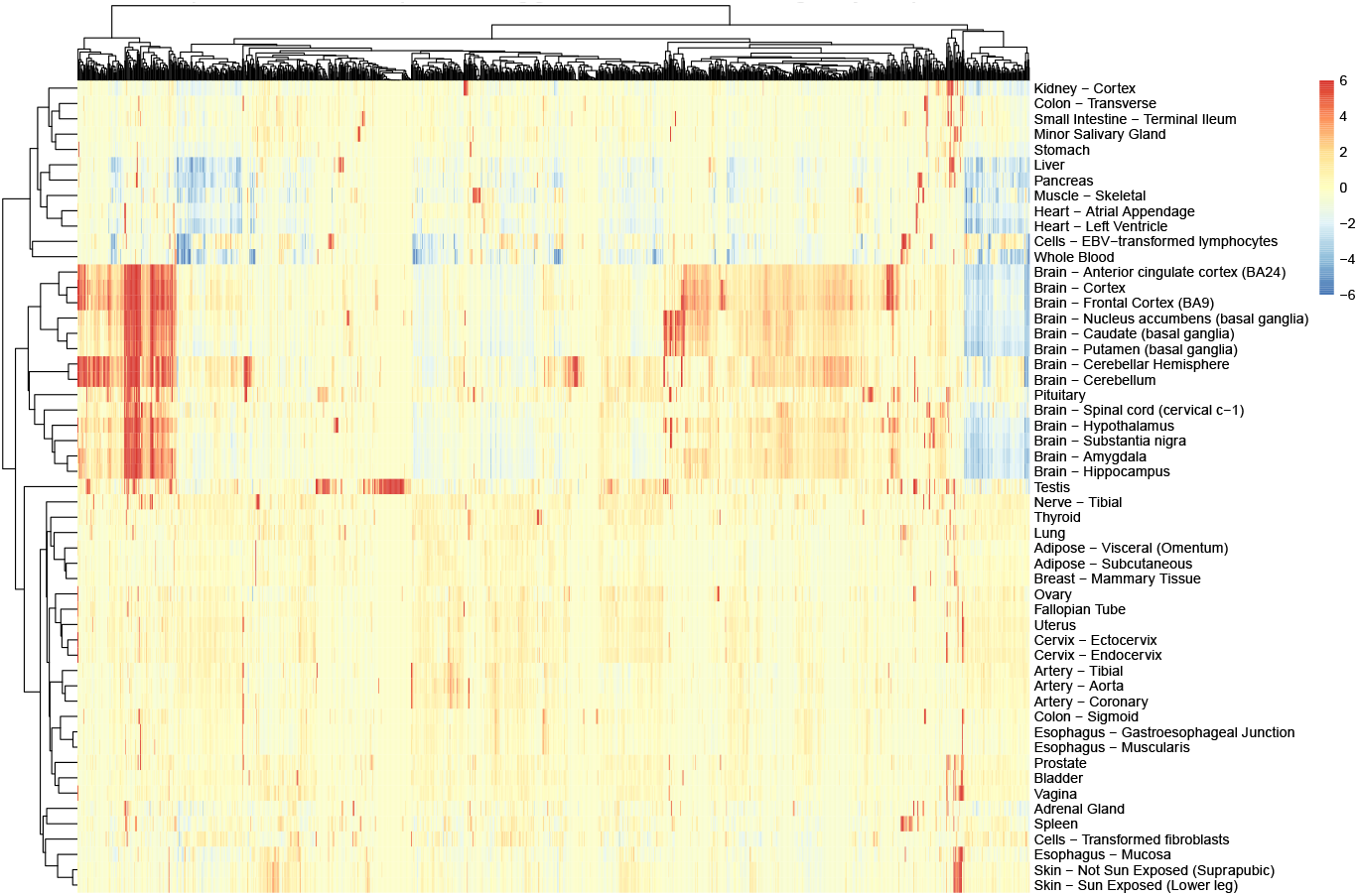
Heatmap of our TS scores of protein coding genes in the GO term of neurological system process across 53 tissues, with rows for tissues and columns for genes. The scores above 6 are truncated to 6 and below −6 to −6.

**Figure 9:**
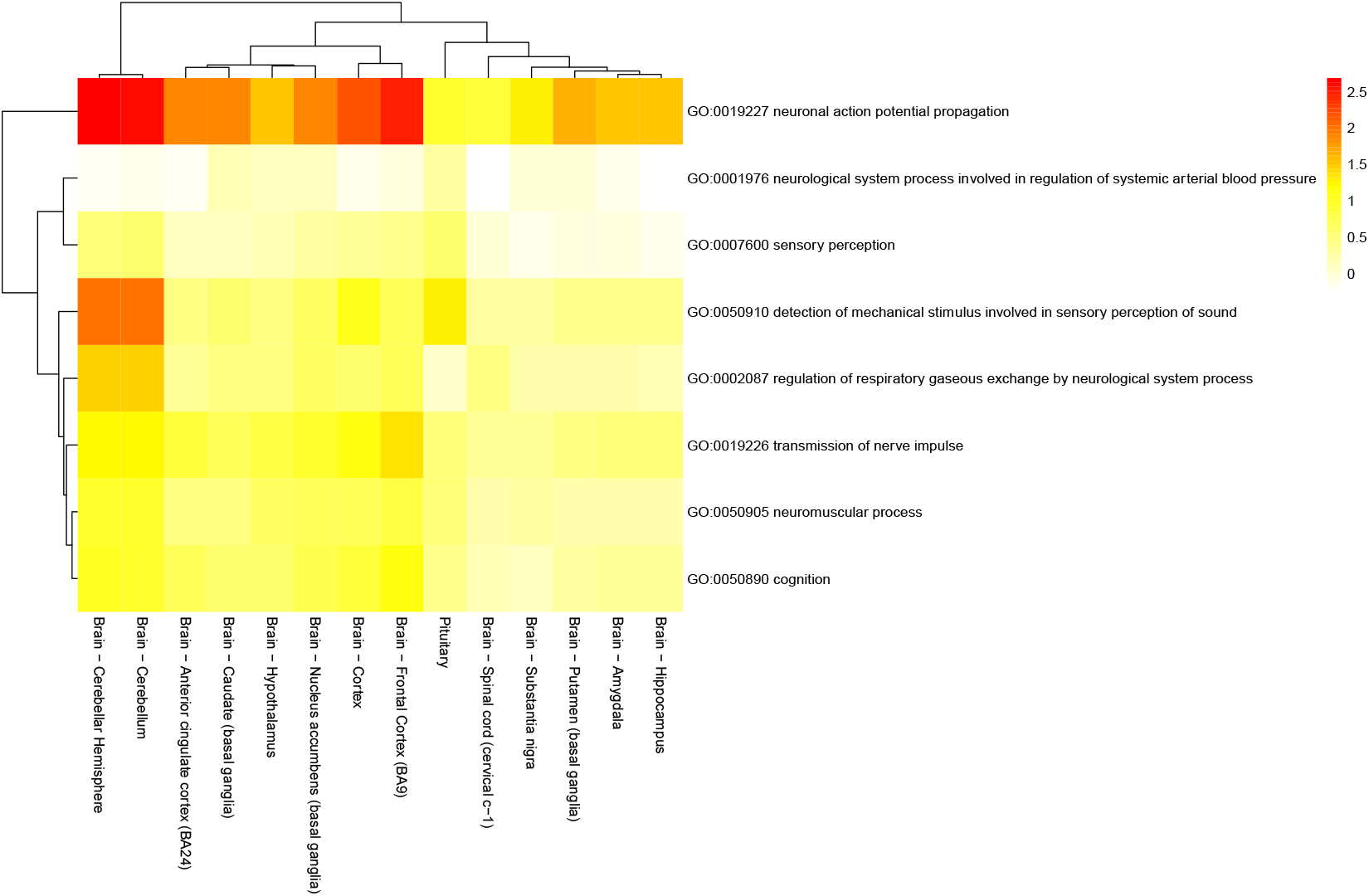
Heatmap of the average of our TS scores from brain related tissues in the children GO terms of the neurological system process GO term.

## 4 Discussion and conclusion

To quantify tissue specificity (TS), our contribution is a new approach, focusing on the population instead of tangling with heterogenous tissue-specific outlier expressions. Our AdaTiSS method is based on our data-adaptive method AdaReg (Wang, et al., 2019), robustly estimating the population distribution and thus constructing the robust and data-adaptive TS scores. Our TS scores quantify the tissue specificity in each tissue for each gene and are comparable across genes. With our quantitative TS cores, it is easy to identify housekeeping genes and tissue enrichment. Our TS scores make it possible to quantitively compare the TS of biology functions such as in the GO term analysis. However, there are still some limitations in the proposed method, so more research is needed.

In the current work, we considered that comparing samples from different tissues, the main effect is the tissue effect, which is confirmed in the explanatory study from sample clustering (Jiang, et al., 2019; Melé, et al., 2015), and thus we modeled the population distribution only including one covariate. There could be other effects such as gender, age affecting the population. One thing needs to notice is that such effects could be associated with the tissue effect, such as some tissue is gender specific. If to remove the gender effect, it may reduce the tissue specificity. If we incorporate more covariates in the population model, there needs careful consideration on the model selection when the samples have outliers. Thus, here we took a simple univariate model on the population and left the problem of model selection on the complex model as a future work.

When considering samples from multiple tissues, we observed the balance inside the population of tissue expressions. The highly expressed outliers may indicate tissue specificity (TS). Therefore, we modeled the population as the symmetric Gaussian distribution. In the supplementary materials, following similar procedures for the Gaussian population, we also developed a data-adaptive procedure for *t*-distribution as the population. We found when the population has heavy tails, our algorithm cannot distinguish which samples in the population tails are outliers, and which are inliers, so the algorithm becomes unstable. Actually, in such cases, the concept of “population” may not be valid. We think it is necessary to predefine the density of the population tails; otherwise all the samples can be in the population. Hence, here we consider the population as Gaussian. As the Gaussian population, there is only one density mode. It could happen there are two modes or even multiple modes, or that the tissue samples might not have any concentrated clusters. To address this concern, based on the current method, we added a diagnosis step: reporting the estimated population proportion. If the proportion is less than 70%, we marked it and checked its fitting. In our future work, we can provide another option for a population with a mixture of multiple Gaussian components. Additional research is also needed to determine the number of components in the population.

Our data-adaptive procedure works best with a light-tailed population such as Gaussian distribution. For the data from the microarray or the mass spectrometry platform, the Gaussian distribution is usually used as a convention after taking the log transformation. For the RNA data from RNA-seq, we worked on the standardized expressions in TPM or RKPM. There are several works analyzing the log of TPM or RKPM for RNA data. In the analysis of transcriptome variations, (Melé, et al., 2015) used the mixed effect model with Gaussian noise, and (Li, et al., 2017) detected the individual outliers within a particular tissue based on the traditional z-score estimated from the normalized log of TPM. In our problem of quantifying TS, we made several adjustments for analyzing the standardized RNA data. First, we took the log transformation to make the expressions more symmetric. As we more focused on the high expressions not on the low expressions, we concentrated the low expression (TPM < 1) at 0 in the log scale to mitigate the inflation from the log transformation. Then, we modified the population concept when the low expressions are not in a small proportion. Finally, we established a criterion to determine whether the low expressions counted for the population.

Another approach of analyzing the RNA expressions is on the counts data based on the negative binomial distribution. (Brechtmann, et al., 2018) developed a method for detecting aberrantly expressed genes using the autoencoder. As they pointed out, their method does not work well if the outlier is not in a small proportion since the autoencoder cannot distinguish the outlier effect from the expected covariates. In our problem where the outliers can be in a large proportion, fitting the negative binomial population given the presence of outliers is more challenging compared to the Gaussian population, since it could be hard to distinguish the over dispersion from the true outliers. There still needs more effort to develop a data-adaptive robust estimation if working on the counts data.

In the analysis pipeline, our proposed method is applied after the preprocessing normalization and/or after removing batch effect. The robustness of the preprocessing steps could affect our TS scores. One approach for future work is to combine the preprocessing steps with quantifying TS to better detect outliers.

Overall, our proposed method provides a robust first step to quantify tissue-specific gene expressions. Our data-adaptive quantitative TS scores (AdaTiSS) bring more precision to functional analysis of tissue samples.

## Acknowledgements

The Genotype-Tissue Expression (GTEx) Project was supported by the Common Fund of the Office of the Director of the National Institutes of Health, and by NCI, NHGRI, NHLBI, NIDA, NIMH, and NINDS. The gene expression data used for the analyses described in this manuscript was obtained from the GTEx Portal in version 7. We acknowledge the discussions with Dr. Hua Tang and Dr. Robert Tibshirani at Stanford.

Conflict of Interest: none declared.

