## Supplemental Material for "AdaTiSS: A Novel Data-Adaptive Robust Method for Quantifying Tissue Specificity Scores"

### Supplementary Materials

**Simulation studies under Gaussian-population.**

In the simulations, we compare the estimation accuracy from various methods under two mixture models: (1) Gaussian mixtures and (2) Gaussian-population mixed with $t$-distribution. For the Gaussian mixtures, consider $X_i \sim iid\ \pi_0 N(\mu_0, \sigma_0^2) + \pi_1 N(\mu_1, \sigma_1^2), i = 1, \dots, n.$ We set $\mu_0 = 0$, $\sigma_0^2 = 1$, and $n = 1000$. Let $\mu_1$be in a sequence $\{0, 1, 3, 5\}$, $\sigma_1^2$ in a sequence $\{0.5^2, 1^2, 1.5^2\}$, and $\pi_1$in a sequence $\{0, 0.1, 0.2, 0.3\}$. To estimate $\mu_0$, the methods under comparisons include the sample mean (when $\gamma = 0$,of our robust methods, labeled by “gam=0”), the sample median (labeled by “med”), the Tukey's biweighting estimation (“biw”), the estimation under the fixed $\gamma = 0.5$ (“gam=0.5”) and fixed $\gamma = 1$ (“gam=1”), our data-adaptive robust estimation under a sequence of $\gamma$'s from 0 to 3 with increment 0.1 (“ada.gam”), the EM algorithm under two Gaussian mixtures (“mix.norm”), and the Efron's local fdr (“locfdr”). We report the mean squared error (MSE) $(\hat{\mu}_0 - \mu_0)^2$ and standard deviation of the squared error for each method under each situation $(\pi_1, \mu_1, \sigma_1)$ from 200 replicates. Since the EM algorithm cannot converge when the null and alternative distributions are not easily to separate, which shows its limitation, we only implement it when $\mu_1 > 1$. To estimate $\sigma_0^2$, since Tukey's biweighting method from the package “affy” does not report the estimation for the variance, we do not take it into comparison. To estimate $\pi_0$, only the $\gamma$ robust estimation methods, the EM algorithm and the local fdr provides the estimation.

Figure S1 - Figure S2 summary the comparison results for estimating $\mu_0$ under Gaussian mixture model, Figure S3 - Figure S4 for estimating $\sigma_0^2$, and Figure S5 - Figure S6 for estimating $\pi_0$. For estimating $\mu_0$, under the case there is no outlier $\pi_1 = 0$, the sample mean (in red bar) shows the smallest error as expected but all other methods under comparison have the error in the similar magnitude. When there exist outliers, the sample mean is no longer robust. Under the case the outlier proportion $\pi_1 = 0.1$ shown in Figure S1, all the robust methods under comparison have similar error while the sample median performs a litter worse. Under the case the large outlier proportion $\pi_1 = 0.3$ shown in Figure S2, our data adaptive robust method shows its advantage: it has much smaller error except in the more undistinguishable case $(\pi_1 = 0.3, \mu_1 = 1, \sigma_1 = 0.5\ or\ 1)$ where our data-adaptive method shows a little large variations. Similar conclusions hold for estimating $\sigma_0^2$ and $\pi_0$.

Under the Gaussian-population mixed with $t$-distribution model, we generate the simulated data from $X_i \sim iid\ \pi_0 N(0,1) + \pi_1 t(\mu_1, df), i = 1, \dots, 1000$, where $\pi_1 \in \{0.1, 0.3\}$, $\mu_1 \in \{1, 3, 5\}$, and $df \in \{3, 5\}$. We report the MSE from 200 replicates in Figure S7 - Figure S9. Since the local fdr method is not designed for the heavy outliers, we do not take it in the comparison in this situation. For the EM algorithm, the issue of unconverging still occurs when $\mu_1 = 1$. Our data-adaptive robust method performs overall good in the various cases.

From our simulation studies, we can see the $\gamma$-robust estimation method is more adaptive to various situations from light outliers to heavy outliers compared to the EM algorithm that only works when the mixtures are well separated and the local fdr is mainly designed under light outliers. Under the selected $\gamma$, our data-adaptive methods perform better overall than the fixed $\gamma$ estimation especially under heavy outliers.

**Data-adaptive procedure embedded in $t$-population mixture model.**

Consider the samples $T_i's$ i.i.d. from a mixture model

$$T_i \sim^{\text{iid}} \pi_0 f_{\nu_0}(t_i) + (1-\pi_0) f_1(t_i) =: f(t_i), \quad i = 1, \ldots, n,$$

where $\pi_0 \in (0.5, 1]$, $f$ is the mixture density function, $f_{\nu_0}$is for the population modeled as $t$-distribution with the degree of freedom $\nu_0$, and $f_1$ is the unknown outlier density. Our goal is under non-vanishing outlier proportion and unknown outlier distribution, to estimate $(\nu_0, \pi_0)$.

Similarly, as in Gaussian-population mixture model, we weigh the mixture model by $f_{\nu_0}^{\gamma}$, where $\gamma \geq 0$, and get

$$f^{(w_0)}(t) = \pi_0^{(w_0)} f_0^{(w_0)}(t) + (1-\pi_0^{(w_0)}) f_1^{(w_0)}(t).$$

We can estimate $f^{(w_0)}$ by

$$w_i = \frac{\frac{1}{n} f_{\nu_0}^{\gamma}(t_i)}{\frac{1}{n}\sum_{j=1}^{n} f_{\nu_0}^{\gamma}(t_j)}.$$

And the theoretical weighted population density is

$$f_0^{(w_0)} = \frac{f_{\nu_0}^{1+\gamma}(t)}{\int f_{\nu_0}^{1+\gamma}(s)\mathrm{d}s} \propto \left(1 + \frac{t^2}{\nu_0}\right)^{\frac{(1+\gamma)(1+\nu_0)}{2}} = \left(1 + \frac{\left(\sqrt{\frac{\nu_0'}{\nu_0}}t\right)^2}{\nu_0'}\right)^{\frac{1+\nu_0'}{2}},$$

which is the density of $\sqrt{\frac{\nu_0}{\nu_0'}}\, T_{\nu_0'}$ , where the degree of freedom for $t$ random variable is $\nu_0' = (1+\nu_0)\gamma + \nu_0$. Under a proper $\gamma$, consider the approximation, $f^{(w_0)} \approx f_0^{(w_0)}$ . The weighted samples $\{(w_i,\ T_i)\}_{i=1}^{n}$ are approximately from purified distribution $\sqrt{\frac{\nu_0}{\nu_0'}}\, T_{\nu_0'}$. Suppose $\nu_0 > 2$ then so is $\nu_0'$. Matching the weighed sample variance to the theoretical variance gives

$$\sum_{i=1}^{n} w_i t_i^2 = \frac{\nu_0}{\nu_0'} \cdot \frac{\nu_0'}{\nu_0' - 2} = \frac{\nu_0}{\nu_0' - 2},$$

shrinking variance by a factor $\frac{\nu_0 - 2}{(\nu_0-2)+(\nu_0+1)\gamma}$. Suppose there is a mean shift $\xi_0$ in the samples, i.e., $T_i - \xi_0 \sim^{iid} f$. Matching the weighted mean to zero gives

$$\sum_{i=1}^{n} w_i (t_i - \xi_0) = 0.$$

Given a fixed $\gamma$, we get

$$\begin{cases} \hat{\xi}_0 = \sum_{i=1}^{n} w_i t_i, \\ \hat{\nu}_0 = \dfrac{(2-\gamma)S^2}{(1+\gamma)S^2 - 1}, \end{cases}$$

where $S^2 = \sum_{i=1}^{n} w_i\,(t_i - \hat{\xi}_0)^2$ . To estimate population proportion, consider unnormalized mixture model and we get

$$\pi_0^{\ddagger} = \frac{\int f(t) f_{\nu_0}^{\gamma}(t) \mathrm{d}t}{\int f_{\nu_0}^{1+\gamma}(t) \mathrm{d}t},$$

$$\hat{\pi}_0 = (\sqrt{\pi \hat{\nu}_0})^{\gamma} \left( \frac{\Gamma(\frac{\hat{\nu}_0}{2})}{\Gamma(\frac{\hat{\nu}_0+1}{2})} \right)^{1+\gamma} \frac{\Gamma(\frac{\hat{\nu}_0+1}{2})}{\Gamma(\frac{\hat{\nu}_0}{2})} \sum_{i=1}^{n} \frac{1}{n} f_{\hat{\nu}_0}^{\gamma}(t_i).$$

To embed the data-adaptive procedure to the $t$-population mixture model, we follow the same procedure as in the Gaussian-population mixture model. For each $\gamma$, we get $(\hat{\nu}_0(\gamma), \hat{\pi}_0(\gamma))$ and estimate the expected fdr under the $t$-population by

$$\hat{E}_{0,t}\widehat{\widetilde{fdr}}(T; \gamma) = \hat{\pi}_0 \sum_{k=1}^{K} \frac{P_{\hat{\nu}_0}(I_k)}{|I_k|/n} \cdot P_{\hat{\nu}_0}(I_k),$$

where $I_k$'s are defined in the same way as in Section 2.3 in the main text. The $\gamma$ is selected by

$$\gamma^* = argmin_{\gamma} |\hat{E}_{0,t}\widehat{\widetilde{fdr}}(T; \gamma) - 1|.$$

Therefore, the data-adaptive estimates are $(\hat{\nu}_0(\gamma^*), \hat{\pi}_0(\gamma^*))$.

We further extend the procedure to scaling $t$-population mixture model. Consider

$$X_i \sim^{iid} (1 - \pi_1)\sigma_0(\mu_0 + t(df = \nu_0)) + \pi_1 f_1.$$

One can first apply $\gamma$-selected procedure under Gaussian-population on $X_i$'s to obtain $(\hat{\mu}_0, \hat{\sigma}_0^2)$, then form $T_i = \frac{X_i - \hat{\mu}_0}{\hat{\sigma}_0}$, next apply $\gamma$-selected procedure under $t$-population on $T_i$'s to obtain $(\hat{\xi}_0, \hat{\tau}_0^2)$, finally obtain the estimate for the population mean for $X_i$'s as $\hat{\mu}_0 + \hat{\xi}_0$ and the estimate for the population variance as $\hat{\sigma}_0^2 \cdot \hat{\tau}_0^2$.

We did a series of simulation studies under $t$-population mixture models with various population degrees of freedom. (We did not show the result plots here.) In our analysis above, we assume $\nu_0 > 2$ to have finite variance, but in practice the algorithm cannot guarantee the estimated $\hat{\nu}_0 > 2$, especially when $\nu_0$ is close to 2, which also indicates the importance and necessity of choosing a proper $\gamma$. However, we found under $\nu_0$ close to 2, the data-adaptive procedure becomes unstable. It is because under heavy tails of population distribution, it is hard to distinguish the samples in the tails to be outliers or inliers from the algorithm. From our experience, to apply our data-adaptive procedure under $t$-population, the population degree of freedom should be at least greater than 5. Therefore, to maintain full power, our data-adaptive procedure prefers light tailed population distribution.

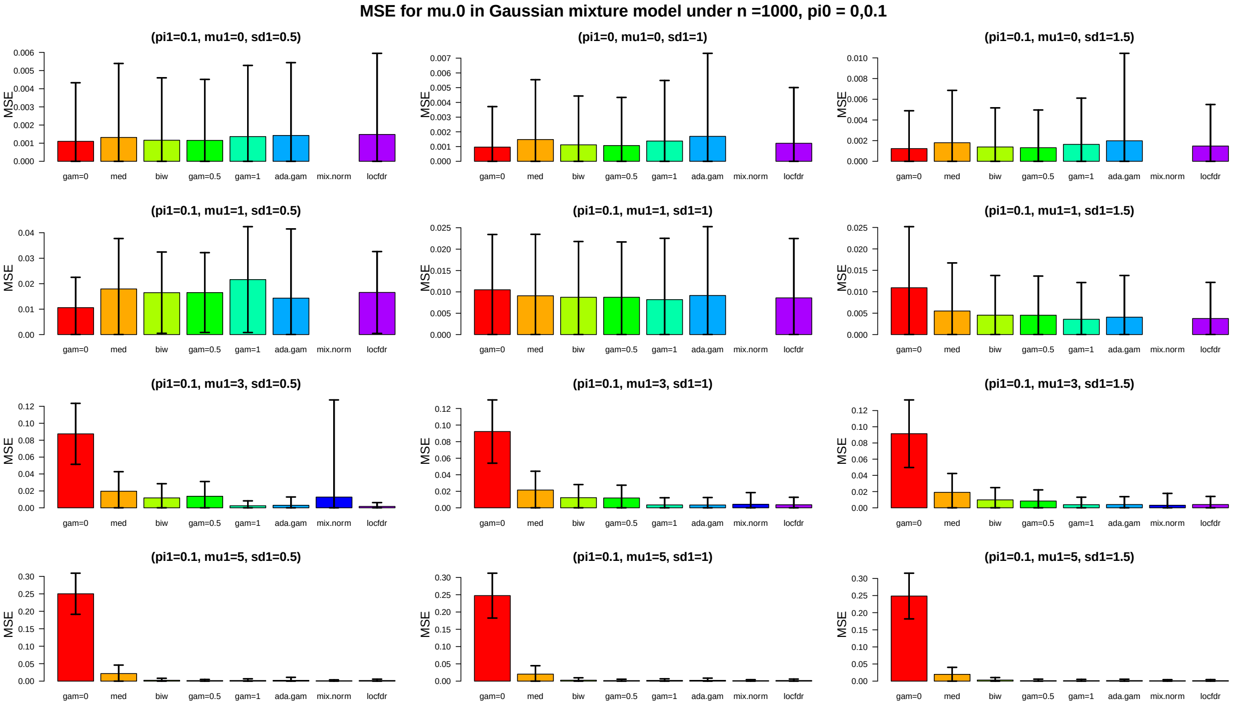


*Figure S1: MSE comparison for estimating $\mu_0$ in the simulated data under 200 independent replicates generated from Gaussian mixture model $X_i \sim iid\ \pi_0 N(0,1) + \pi_1 N(\mu_1, \sigma_1^2), i = 1, \dots, 1000$ where $\pi_1 = 0, 0.1,\ \mu_1 \in \{0, 1, 3, 5\}$ and $\sigma_1^2 \in \{0.5^2, 1^2, 1.5^2\}$, The error bars are based on 2 times standard deviation of the squared error from 200 replicates.*

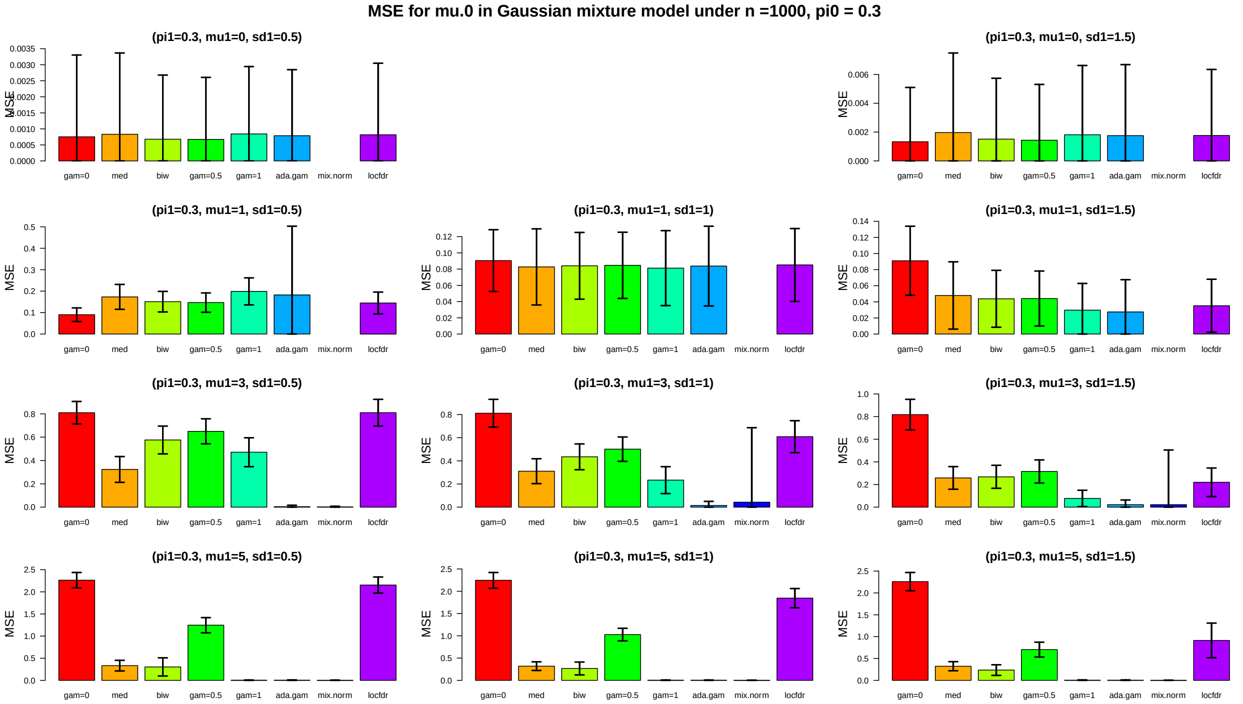


*Figure S2: The same caption as in Figure 1 except that $\pi_1 = 0.3$ in the generating model.*

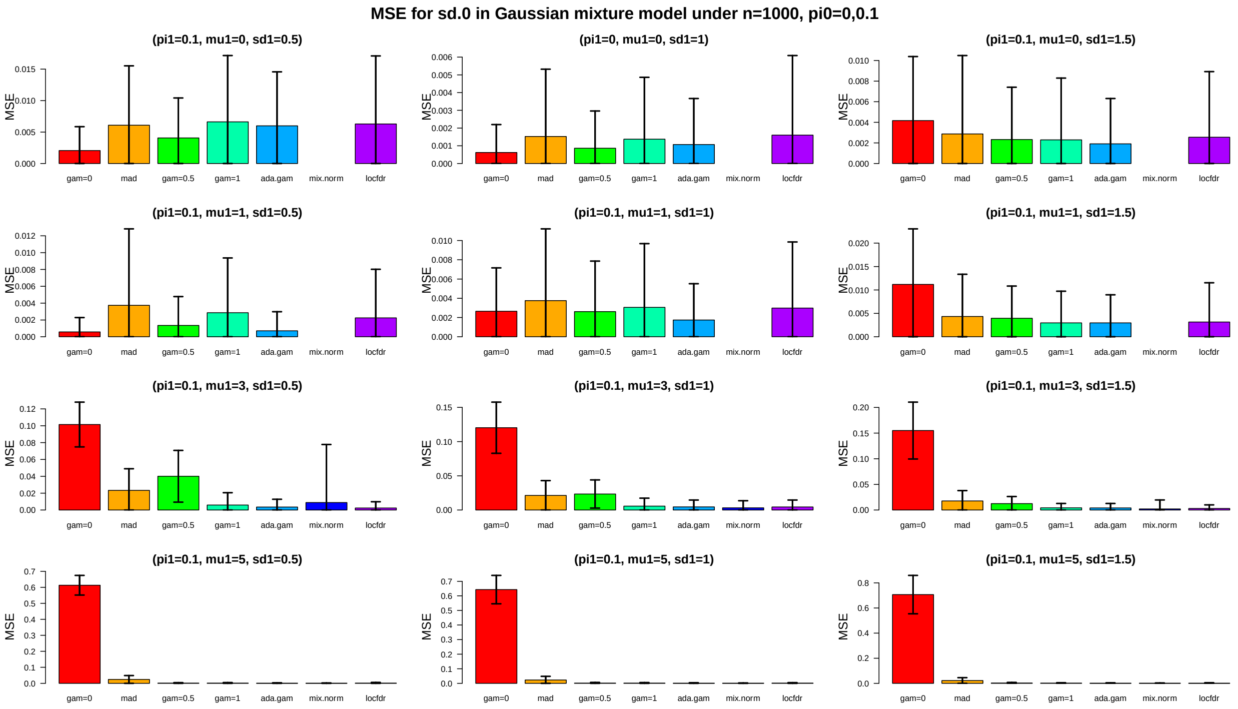


*Figure S3: MSE comparison for estimating $\sigma_0^2$ in the simulated data under 200 independent replicates generated from Gaussian mixture model $X_i \sim iid\ \pi_0 N(0,1) + \pi_1 N(\mu_1, \sigma_1^2), i = 1, \ldots, 1000$ where $\pi_1 = 0, 0.1,\ \mu_1 \in \{0,1,3,5\}$ and $\sigma_1^2 \in \{0.5^2, 1^2, 1.5^2\}$, The error bars are based on 2 times standard deviation of the squared error from 200 replicates.*

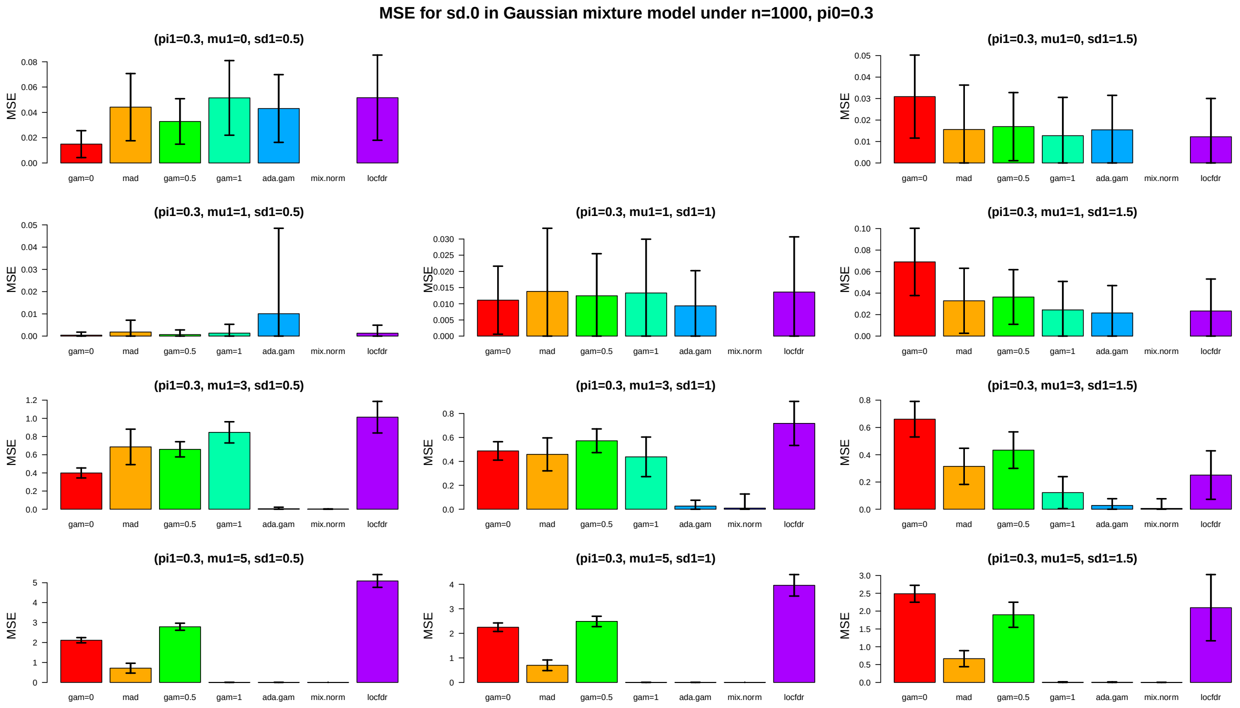


*Figure S4: The same caption as in Figure 3 except that $\pi_1 = 0.3$ in the generating model.*

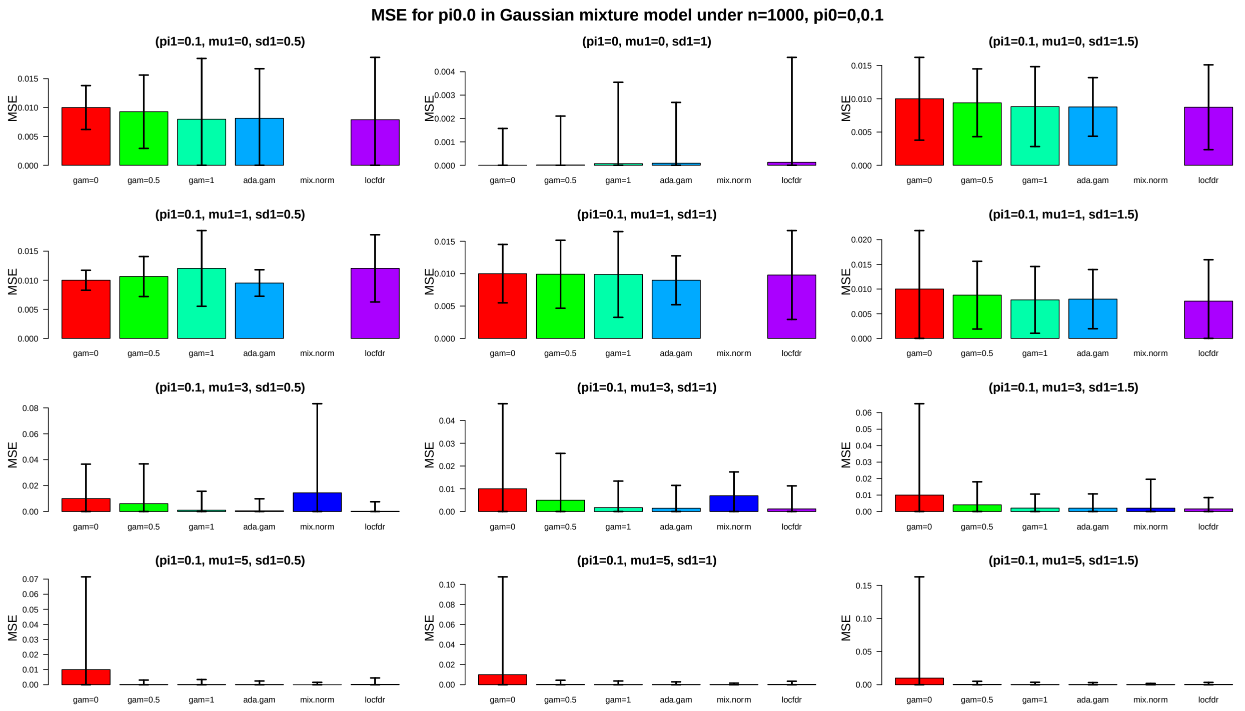


*Figure S5: MSE comparison for estimating $\pi_0$ in the simulated data under 200 independent replicates generated from Gaussian mixture model $X_i \sim iid\ \pi_0 N(0,1) + \pi_1 N(\mu_1, \sigma_1^2), i = 1, \ldots, 1000$ where $\pi_1 = 0{,}0.1$, $\mu_1 \in \{0,1,3,5\}$ and $\sigma_1^2 \in \{0.5^2, 1^2, 1.5^2\}$, The error bars are based on 2 times standard deviation of the squared error from 200 replicates.*

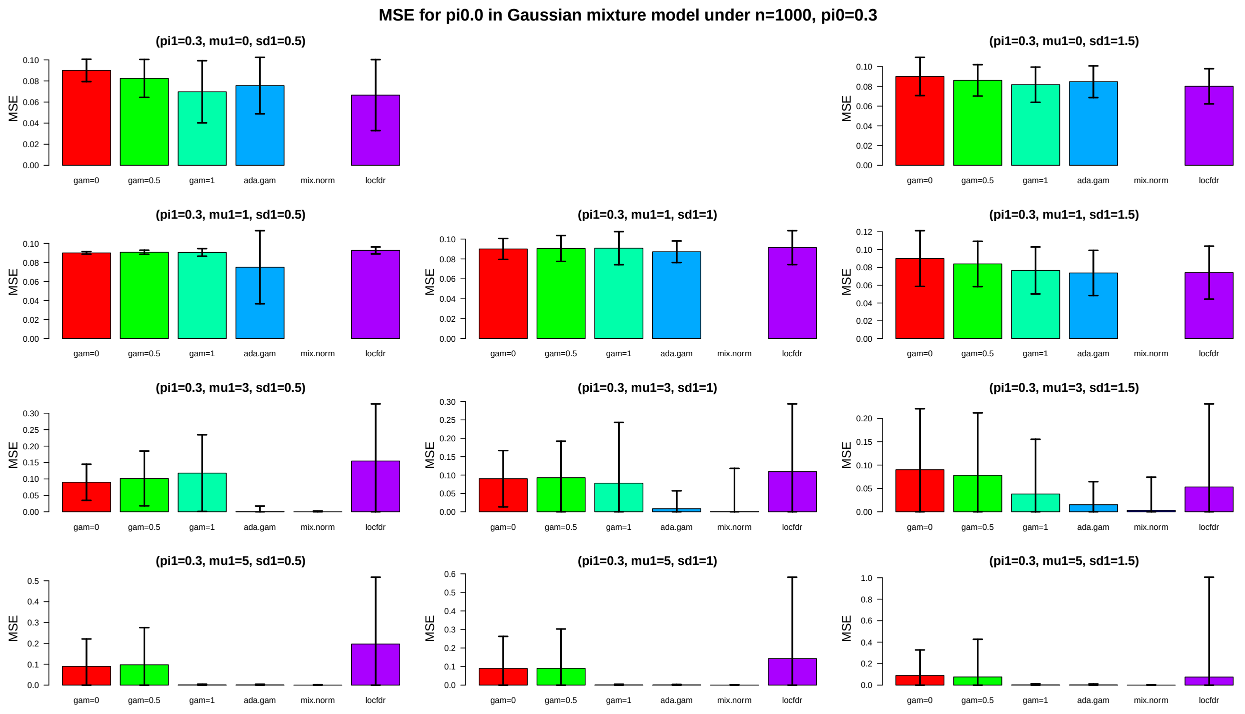


*Figure S6: The same caption as in Figure 5 except that $\pi_1 = 0.3$ in the generating model.*

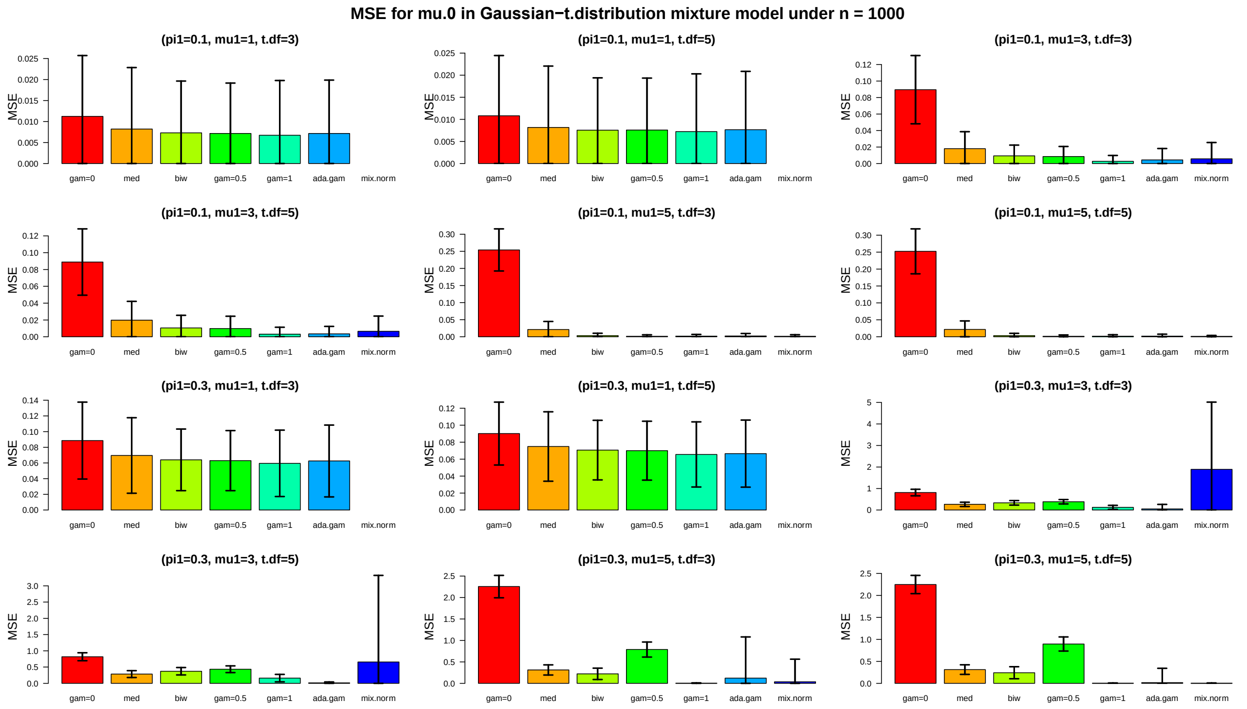


*Figure S7: MSE comparison for estimating $\mu_0$ in the simulated data under 200 independent replicates generated from Gaussian and t-distribution mixture model $X_i \sim iid\ \pi_0 N(0,1) + \pi_1 t(\mu_1, df), i = 1, \dots, 1000$, where $\pi_1 \in \{0.1, 0.3\}$, $\mu_1 \in \{1, 3, 5\}$, and $df \in \{3, 5\}$. The error bars are based on 2 times standard deviation of the squared error from 200 replicates.*

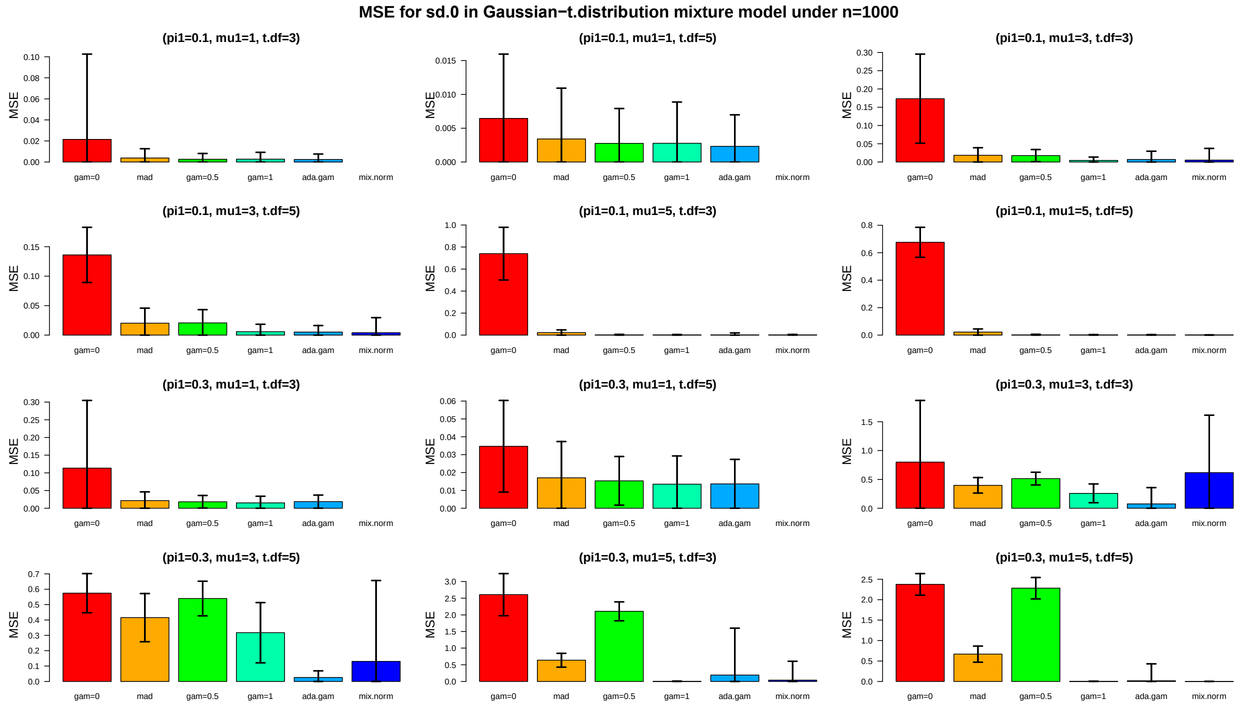


*Figure S8: The same caption as in Figure 7 except the comparison for estimating $\sigma_0^2$.*

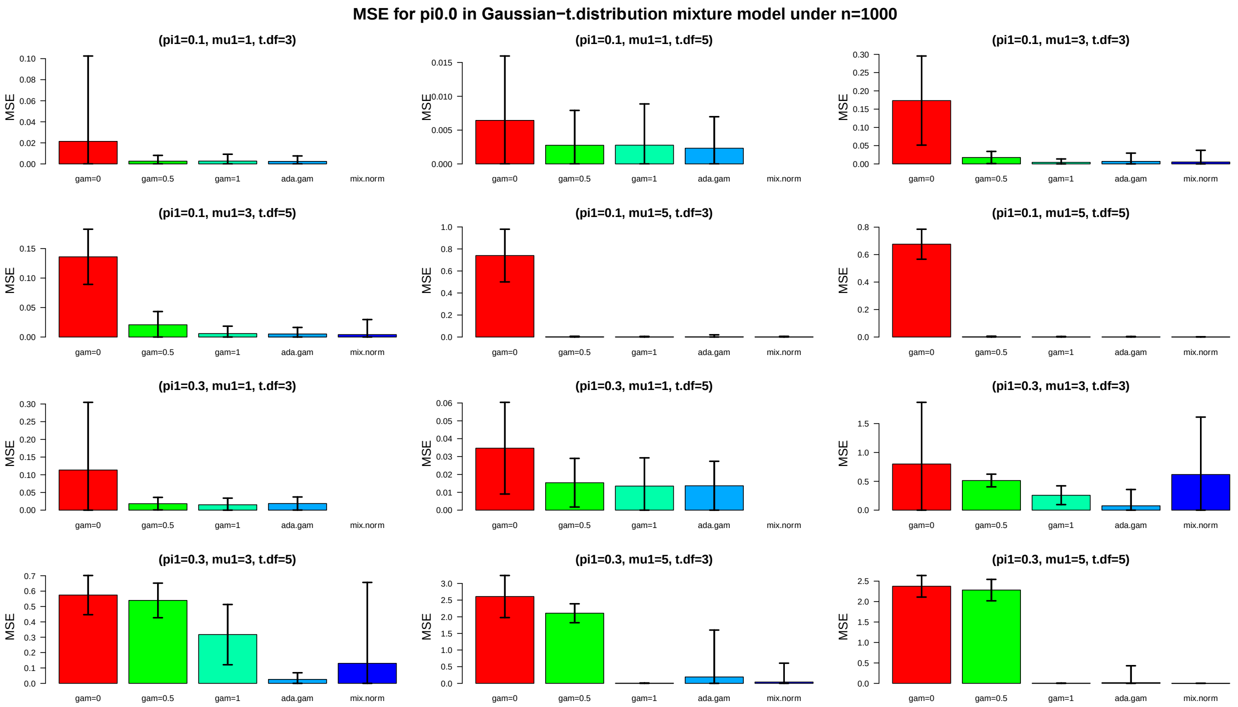


*Figure S9: The same caption as in Figure 7 except the comparison for estimating $\pi_0$.*
